# Proteolytic activation of diverse antiviral defense modules in prokaryotes

**DOI:** 10.1101/2025.11.14.688446

**Authors:** Simone A. Evans, Jianxiu Zhang, Yannie Lam, Katherine Wang, Yu Tin Lin, Daniel Herschlag, Liang Feng, Alex Gao

## Abstract

Linked protease–effector modules are widespread in prokaryotic antiviral defense, yet the mechanisms of most remain poorly understood. Here we show that four of the most prevalent modules—metallo-β-lactamase (MBL)-fold hydrolase, α/β-hydrolase, Pepco, and EACC1—form latent death effectors that are unleashed by site-specific proteolysis. Genetic, biochemical, and structural analyses reveal novel modes of effector licensing. MBL and α/β-hydrolase are zymogens activated by cleavage at two distinct sites, and upon proteolysis, MBL becomes a Zn^2+^-dependent double-stranded DNA nuclease. In contrast, Pepco and EACC1 act as pore-forming toxins via distinct mechanisms: Pepco constitutively oligomerizes into a denaturation-resistant beta barrel that is activated by cleavage after a specific isoleucine in its C-terminal tail, whereas EACC1 monomers assemble into large membrane pores following removal of an autoinhibitory domain. All four modules are fused to diverse sensors to detect a wide range of phage signals, and EACC1–DnaK chaperone fusions suggest a convergence between defense and general stress responses. These findings establish proteolysis-gated activation as a dominant, modular logic for anti-phage defense and reveal parallels with eukaryotic innate immunity.

## Introduction

Evolutionary pressures exerted by bacteriophages drive the evolution of diverse antiphage immune systems in prokaryotes^1–3^. These immune systems frequently have a modular architecture, with diverse sensors and effectors serving as interchangeable parts that can be reshuffled and transferred horizontally across species^4–7^. This modularity facilitates rapid innovation through new sensor–effector pairings, potentially improving detection and neutralization of diverse phage threats.

Protease domains are a recurrent feature of modular immune architectures in prokaryotes, often encoded next to proteins with unknown functions. Recent computational studies have highlighted the abundance of protease-associated proteins across diverse defense contexts, suggesting a functional linkage between proteases and antiviral defense systems^4–8^. These include the small hypothetical proteins Pepco (also called Trypco1 and Trypco2)^6,8^ and EACC1 (ref.^6,7^), as well as a metallo-β-lactamase (MBL)-like enzyme^4,5,9^, an α/βfold hydrolase^6,7,10^, caspase associated regulator component (CATASP)^8,10^, and a metallophosphoesterase (MPE)-like enzyme^9^. While a few defense-associated protease systems have been characterized, including the CRISPR-associated sigma factor Csx30 (ref.^11–13^), CRISPR-associated antisigma factor CalpT^14^, bacterial gasdermin^15,16^, and caspase-like prokaryotic proteases (PCaspase)^17^, the functions of many widespread modules and their integration into defense systems remain poorly understood.

We characterize the activation mechanisms of four of the most broadly distributed protease-associated effectors—MBL, α/β hydrolase, Pepco, and EACC1—revealing a conserved logic in which site-specific proteolytic cleavage activates latent death effectors. We find that MBL and α/β hydrolase are both inactive zymogens that are cleaved into three fragments to induce cell death. MBL is a double-stranded DNA (dsDNA) nuclease with an active site occluded by linkers that are removed by cleavage. In contrast, Pepco and EACC1 are pore-forming toxins that are activated by a single cleavage event per monomer. Pepco constitutively forms a multimeric, denaturation-resistant βbarrel that is activated by removal of its C-terminal tail, whereas EACC1 is an autoinhibited monomer that assembles into a large membrane pore after cleavage. Pepco barrels resemble miniature versions of the membrane attack complex/perforin (MACPF)^18^ found in mammalian immune systems and pore-forming toxins of pathogenic bacteria^19,20^. All four of these protease–effector pairs are coupled with structurally diverse sensor domains across bacterial taxa, implying the ability to respond to various distinct infection-induced signals. Our findings demonstrate that across distant lineages and diverse effectors, protease-mediated autoinhibition is a recurring strategy to maintain lethal proteins silent until they are needed.

### Protease–effector modules are widespread in prokaryotes and associated with diverse sensors

To assess the prevalence and distribution of protease–effector modules in prokaryotes, we performed PSI-BLAST searches against the NCBI non-redundant protein sequence database (nr) to identify homologs of all known and predicted modules. Across the ten surveyed families^6-17^, abundance varied by nearly three orders of magnitude, ranging from the rare Csx30 to the widespread Pepco (Fig. 1A). Apart from PCaspase^17^, four of the five most abundant modules remain uncharacterized: MBL, α/β hydrolase, Pepco, and EACC1. These effectors are broadly distributed across bacterial phyla, including multicellular taxa such as cyanobacteria and actinomycetes. We therefore sought to investigate their functional relationship with their associated proteases.

**Figure 1.**
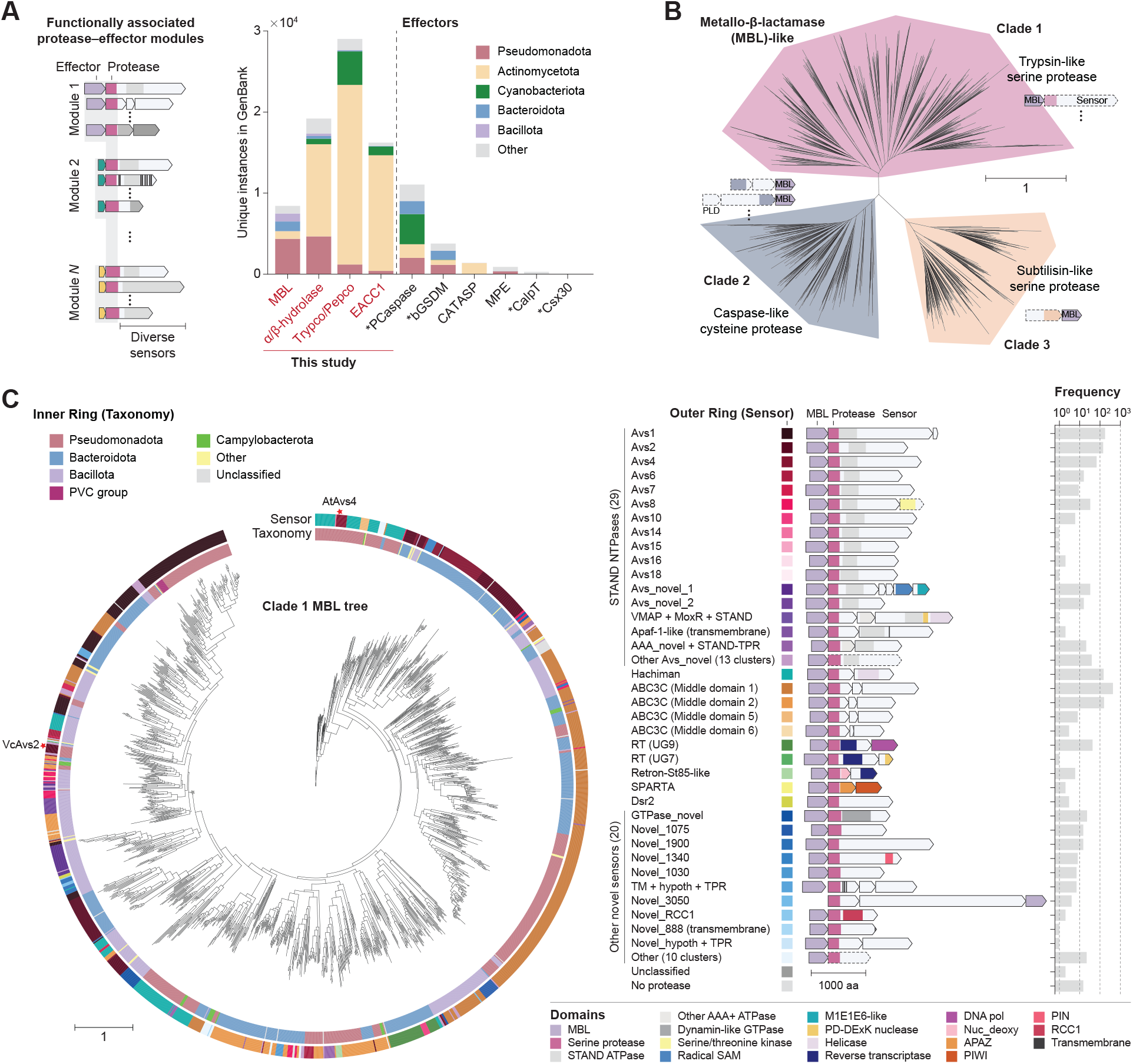
Protease-associated effector domains in prokaryotes. (**A**) Number of unique instances of protease-associated effector domains. Abbreviations: MBL, metallo-β-lactamase; Pepco, peptidase co-occurring domain; EACC1, Effector-Associated Constant Component 1; PCaspase, prokaryotic caspase; bGSDM, bacterial gasdermin; CATASP, CATRA-associated small protein; MPE, metallophosphoesterase; CalpT, CRISPR-associated Lon-SAVED protease target; Csx30, CRISPR-associated accessory protein. Asterisk (^*^) indicates domains previously investigated experimentally. (**B**) Maximum likelihood tree of protease-associated MBL representatives clustered at 95% sequence identity, encoded adjacent to trypsin-like serine proteases (Clade 1), caspase-like proteases (Clade 2), and subtilisin-like serine proteases (Clade 3). (**C**) Maximum likelihood tree of MBL Clade 1. The inner ring denotes host taxonomy, and the outer ring indicates the immune sensor associated with the protease domain. Representative domain architectures and the frequency of each protease–MBL system in GenBank are shown. Stars in the outer ring mark homologs experimentally investigated in this study.

We first focused on protease-associated MBL-fold hydrolases across diverse bacterial lineages. Phylogenetic analysis revealed three major clades, each associated with distinct protease classes: trypsin-like serine proteases, caspase-like cysteine proteases, and S8 subtilisin-like proteases (Fig. 1A). Comprehensive analysis of the serine protease clade demonstrated phylogenetic congruence between the MBL and protease, suggesting co-evolution and functional inter-dependence of these two components (Supplementary Fig. 1). Moreover, these modules are fused to a remarkable diversity of putative sensor domains, encompassing 29 distinct STAND (Signal Transduction ATPases with Numerous Domains) families^4,5,21,22^, Hachiman^2^, four types of ABC3C systems^9^, two distinct defense-associated reverse transcriptases (DRTs)^23^, an St85-like retron, SPARTA^24^, Dsr2 (ref.^4^), and 20 uncharacterized types (Fig. 1E and Supplementary Fig. 2). Additional, distinct sensor associations are found in the cysteine and subtilisin-like protease clades (Fig. 1B). Among the STAND families are sensors for phage proteins, including the large terminase (Avs1-2) and portal (Avs4)^5^. In addition, Hachiman senses genome integrity^25^ while SPARTA senses phage ssDNA^24^. Together, these findings suggest that MBL–protease modules can respond to a broad range of phage-derived input signals during defense.

**Figure 2.**
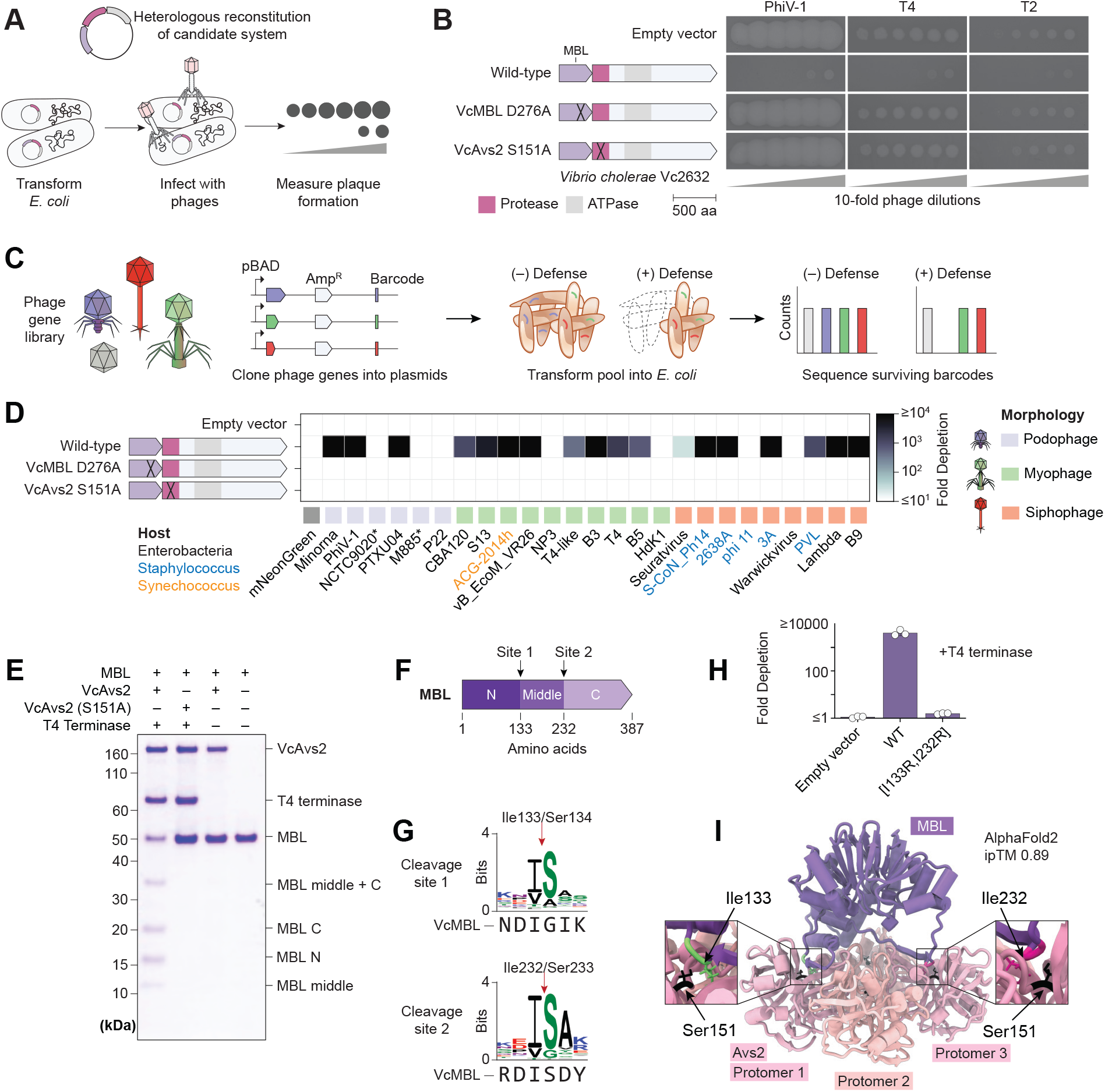
The MBL zymogen is cleaved at two distinct sites by its associated serine protease. (**A**) Schematic of phage plaque assay. (**B**) Heterologous reconstitution of the *Vibrio cholerae* protease (VcAvs2) and its associated VcMBL effector in *E. coli*. Plaque assays show 10-fold serial dilutions of phages PhiV-1, T4, and T2 on strains expressing wild-type or catalytic mutants of VcMBL (D276A) and VcAvs2 protease (S151A). (**C**) Schematic of the plasmid depletion assay library design. (**D**) Co-expression toxicity assay in *E. coli* showing the VcMBL–protease defense system or its catalytic mutants expressed with diverse phage terminase proteins. (**E**) SDS-PAGE analysis of VcMBL cleavage by VcAvs2. (**F**) Cleavage sites of VcMBL (I133 and I233), identified by Edman degradation (see Fig. S3). (**G**) Sequence logo of the VcMBL cleavage motif. (**H**) Co-expression toxicity assay in *E. coli* with T4 terminase and the VcMBL–protease defense system or VcMBL cleavage site mutants (I133R, I233R). (**I**) AlphaFold2 multimer model of VcMBL in complex with a C4-symmetric VcAvs2 protease tetramer, showing catalytic residues positioned near the predicted cleavage sites on VcMBL.

### Phage-activated protease cleaves MBL at two distinct sites

To investigate the function of MBL–protease modules, we focused on homologs within antiviral STAND (Avs) families, which are activated by phage proteins^5,21^, enabling reconstitution of defense activity in the absence of phage. We cloned two MBL homologs, one associated with Avs2 from *Vibrio cholerae* Vc2632 (VcAvs2) and another associated with Avs4 from *Acinetobacter towneri* A6 (AtAvs4). Avs2 and Avs4 recognize monomers of the phage large terminase subunit and portal, respectively, which are core components of the conserved DNA packaging machinery of tailed phages.

Heterologous expression of VcMBL and VcAvs2 in *E. coli* conferred defense against phages PhiV-1, T4, and T2 in plaque assays (Fig. 2A–B). Co-expression of the system with terminase proteins triggered cellular toxicity (Fig. 2C–D), consistent with defense activation^5^. Catalytic mutations in either VcMBL (D276A) or the VcAvs2 protease (S151A) abolished both phenotypes (Fig. 2B,D). Reconstitution of AtMBL and AtAvs4 in *E. coli* similarly conferred defense against PhiV-1 and exhibited phage portal-dependent toxicity (Fig. 3A–B). These results demonstrate that both MBL and its associated protease are required for defense.

**Figure 3.**
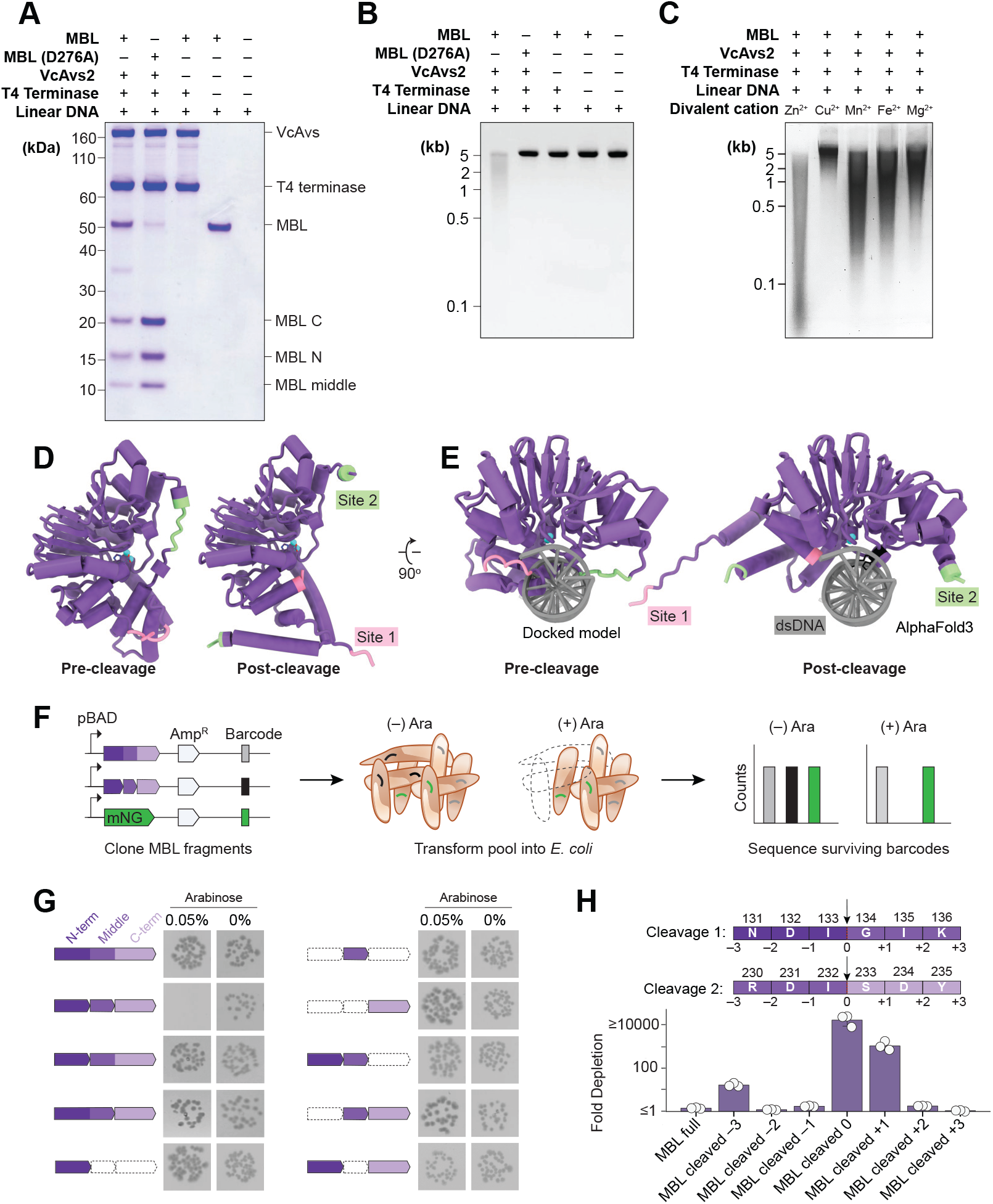
MBL degrades dsDNA and requires all three cleaved fragments for activity. (**A**) SDS-PAGE gel of in vitro cleavage of wild-type VcMBL or catalytically inactive VcMBL (D276A) by VcAvs2. (**B**) Agarose gel showing degradation of linear dsDNA by cleaved VcMBL in the presence of Zn^2^+. (**C**) Agarose gel of VcMBL dsDNAse activity in the presence of different divalent metal cations. (**D**) Comparison of AlphaFold3 models of full-length and cleaved VcMBL. Cleavage site 1 is shown in pink; cleavage site 2 in green. (**E**) Cleaved VcMBL model is co-folded with DNA. Full-length VcMBL is aligned to DNA from the cleaved VcMBL–DNA complex. (**F**) Schematic of pooled VcMBL fragment library design. (**G**) Expression toxicity of VcMBL fragment combinations in *E. coli* to mimic proteolytic cleavage. (**H**) Expression toxicity in *E. coli* of VcMBL fragments with shifted cleavage sites.

To test for direct interactions between MBL and protease, we purified VcMBL, VcAvs2, and T4 terminase as recombinant proteins in *E. coli* and incubated them *in vitro* in the presence of ATP and Mg^2+^. VcMBL was cleaved into three distinct fragments, and cleavage required catalytically active protease and the presence of T4 terminase (Fig. 2E). Edman degradation mapped the cleavage sites to Ile133/Ser134 and Ile232/Ser233 (Supplementary Fig. 4). The presence of a single partial cleavage product, corresponding to the middle and C-terminal fragments, suggested that the first site is cleaved more efficiently than the second. Sequence analysis of 420 homologs revealed partial conservation of the residues immediately flanking the cleavage site (Fig. 2G). Mutating both –1 (P1) residues (I133R/I232R) eliminated cellular toxicity (Fig. 2H), whereas mutating the +1 (P1’) residues had no effect on VcMBL but partially reduced AtMBL activity (Supplementary Fig. 5A–B). In contrast, mutations at less conserved positions near the cleavage site in both VcMBL and AtMBL did not affect activity (Supplementary Fig. 5A–B).

Because Avs2 tetramerizes upon terminase binding, we modeled VcAvs2 and AtAvs4 protease domain tetramers in complex with their cognate MBLs using AlphaFold2 multimer (ipTM 0.89 for both models). In both cases, the catalytic serines of two distinct protomers aligned with the MBL cleavage sites (Fig. 2I and Supplementary Fig. 3C). Notably, VcAvs2 is predicted to employ diagonal protomers of the tetramer to cleave MBL, whereas AtAvs4 uses adjacent protomers (Fig. 2I and Supplementary Fig. 3C,E–F). Disruption of as many as three predicted charge interactions and β-strand contacts at the MBL–protease interface did not abolish toxicity (Supplementary Fig. 5C–D), suggesting that multiple, partially redundant interactions may collectively stabilize the interface. Together, these structural models indicate that protease tetramerization upon phage recognition may help position the catalytic residues for efficient double cleavage of MBL.

### Cleaved MBL is a Zn^2+^-dependent dsDNA nuclease

Because protease-associated MBLs share remote homology with the MBL-superfamily nucleases ComEC and RNaseJ^26^, we tested whether VcMBL exhibits nuclease activity. Cleaved VcMBL (100 nM) degraded both linear and circular dsDNA in the presence of Zn^2+^ (Fig. 3A–B and Supplementary Fig. 6A), but did not cleave RNA or ssDNA (Supplementary Fig. 6B). Notably, a catalytically inactive MBL mutant (D276A) was cleaved but failed to degrade dsDNA, suggesting that proteolytic cleavage and nuclease activity are separate processes. Metal ion substitution assays showed robust activity with Zn^2+^, reduced activity with Mn^2+^, Fe^2+^, or Mg^2+^, and minimal activity with Cu^2+^ (Fig. 3C). These results are consistent with the active site not accommodating the square-planar coordination of Cu^2+^, in contrast to the tetrahedral geometries of other divalent cations. Thus, VcMBL appears to prefer Zn^2+^, in line with other MBL superfamily members.

Structural modeling with AlphaFold3 suggested that two linker regions of full VcMBL occlude the active site, preventing dsDNA access (Fig. 3D–E). Cleavage of these linkers exposes the active site, activating MBL as a nonspecific nuclease. Models of cleaved AtMBL predicted similar dsDNA engagement and linker-dependent constraints (Supplementary Fig. 6C–D), indicating that cleavage of inhibitory linkers could be a conserved mechanism of MBL activation by serine proteases.

To define the minimal components required for dsDNA cleavage, we expressed VcMBL fragments individually and in combinations under an inducible promoter (Fig. 3F). Only the set of all three VcMBL fragments induced cellular toxicity, whereas any single fragment or fragment pair did not (Fig. 3G). Likewise, a point mutation to each fragment of AtMBL ablated cellular toxicity (Supplementary Fig. 3D). Toxicity required the fragment junctions to be at or immediately downstream (+1) of the scissile bonds. In contrast, no activity was observed when the junctions were placed upstream (1 to 3) or further downstream (+2, +3) (Fig. 3H). These results suggest that accurate protease cleavage is essential for activity.

### Double cleavage of an α/β-hydrolase zymogen induces cell death

A predicted α/β-fold hydrolase of unknown function is widespread in Actinobacteria and Pseudomonadota (formerly Proteobacteria) (Fig. 1A) and associated with proteases that are fused to diverse sensor domains, including β-transducin repeats and an uncharacterized STAND NTPase^27^ linked to type I-E CRISPR systems (Fig. 4A). We investigated a representative from the soil bacterium *Lysobacter capsici* AZ78, comprising an α/βhydrolase (LcHydrolase) and a caspase-like cysteine protease (LcCaspase). We purified both proteins recombinantly from *E. coli* and found that LcCaspase progressively cleaved LcHydrolase at two sites, and that cleavage required a catalytically active protease (Fig. 4B). The resulting N-terminal (8.1 kDa) and C-terminal (19.8 kDa) fragments were readily detected by SDS–PAGE. The small middle fragment (4.7 kDa) was not visible due to its size and low numbers of Coomassie-stainable residues, but was inferred through partial cleavage intermediates observed at early time points (Fig. 4B). The putative cleavage sites of LcHydrolase, Arg73 and Arg117, are positioned adjacent to the catalytic residue (Cys129) of two LcCaspase protomers in an AlphaFold3 model of the complex (ipTM 0.91) (Fig. 4C–D).

**Figure 4.**
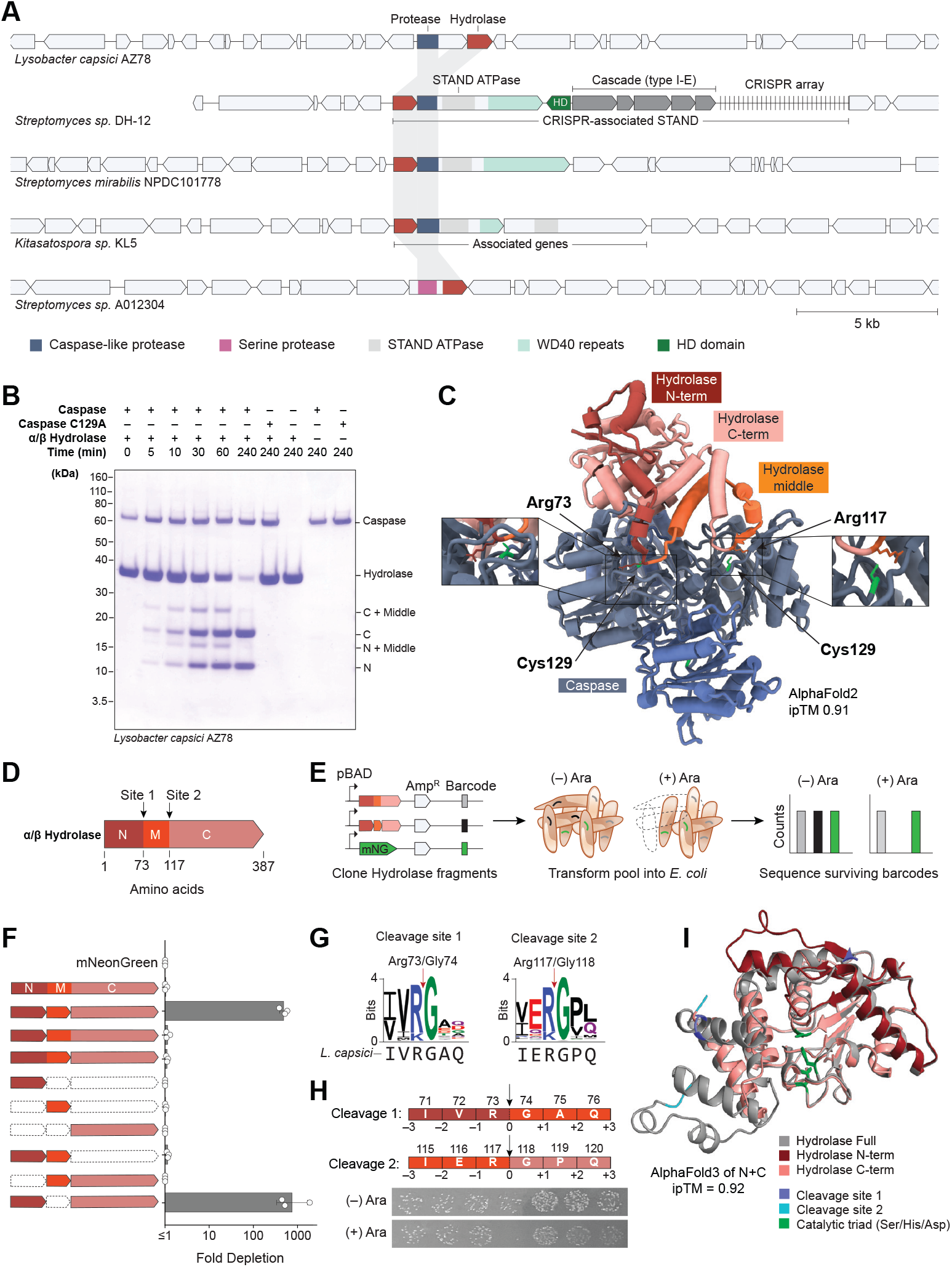
α/β-hydrolase zymogen is cleaved at two distinct sites by its associated protease. (**A**) Representative genomic neighborhoods of α/β-hydrolase–protease pairs. (**B**) Time-course assay of *Lysobacter capsisi* hydrolase (LcHydrolase) cleavage by LcCaspase. (**C**) AlphaFold3 model of LcHydrolase in complex with an LcCaspase tetramer (ipTM 0.91), with catalytic residues of diagonal caspase protomers aligned at the predicted cleavage sites on LcHydrolase. (**D**) Schematic of the putative cleavage sites of LcHydrolase. (**E**) Schematic of the LcHydrolase fragment library design for toxicity assay. (**F**) Expression toxicity assay of LcHydrolase fragment combinations in *E. coli* to mimic proteolytic cleavage. (**G**) Sequence logo of LcHydrolase cleavage sites. (**H**) Expression toxicity of LcHydrolase fragments with shifted cleavage sites. (**I**) AlphaFold3 model of full LcHydrolase (gray) aligned with a model of cleaved LcHydrolase (red and coral).

To assess the functional consequence of cleavage, we expressed LcHydrolase fragments individually and in combinations under an inducible promoter in *E. coli* (Fig. 4E). Expression of all three fragments, or just the N- and C-terminal fragments, triggered cell death, whereas the individual fragments alone did not (Fig. 4F). These results suggest that LcHydrolase functions as a protease-activated death effector downstream of LcCaspase. Analysis of 161 homologs revealed conserved cleavage motifs at Arg73 and Arg117, where arginine is followed by glycine (R↓G) (Fig. 4G). Cellular toxicity required the fragment junctions to be precisely at the scissile bond, as no activity was observed when the junctions were placed –3 to +3 residues upstream or downstream (Fig. 4H).

Structural modeling of the cleaved two-fragment state (ipTM 0.92) suggested conformational rearrangements relative to intact LcHydrolase. Cleavage within the helix spanning residues Asp64–Arg73 may permit its terminal segment to refold into a β-strand that extends the core hydrolase β-sheet (Fig. 4I and Supplementary Fig. 7A–D). The N terminus of the C-terminal fragment contributes to the putative “lid” region of the hydrolase^28,29^, and removal of the central fragment after the second cleavage event may allow this region to adopt its correct position, enabling proper lid formation.

Unlike MBL, cleaved LcHydrolase did not degrade nucleic acids (Supplementary Fig. 8A–C). HHpred^30^ analysis revealed closest homology to lipases and oligopeptidases, suggesting possible activity on ester-or peptide-containing substrates. Further analysis will be required to identify the substrate of the hydrolase.

### Pepco forms stable β-barrels that are proteolytically activated by C-terminal cleavage

Pepco is a small protein (117 amino acids) with no known enzymatic activity that is associated with serine or cysteine proteases and prevalent in Actinobacteria and cyanobacteria^6,8^ (Fig. 1A and 5A). Associated proteases are fused to diverse sensors, including formylglycine-generating enzymes (FGE), transmembrane STAND ATPases, and peptidoglycan-binding domains (Fig. 5A). We focused on a representative from *Streptomyces subrutilus* 10-1-1, where Pepco (SsPepco) is encoded immediately upstream of an Avs4 homolog (SsAvs4) containing a trypsin-like serine protease domain. Co-expression of SsPepco and SsAvs4 with phage portals in *E. coli* triggered strong cellular toxicity, consistent with Avs4 activation (Fig. 5B). Toxicity was abolished by deletion of SsPepco or mutation of the protease catalytic serine (S151A). However, replacing the SsPepco–protease pair with a PD-DExK nuclease domain from a related *Streptomyces exfoliatus* Avs4 homolog restored portal-dependent toxicity (Fig. 5B), suggesting that SsPepco functions as a modular component that can be exchanged between diverse effector domains.

**Figure 5.**
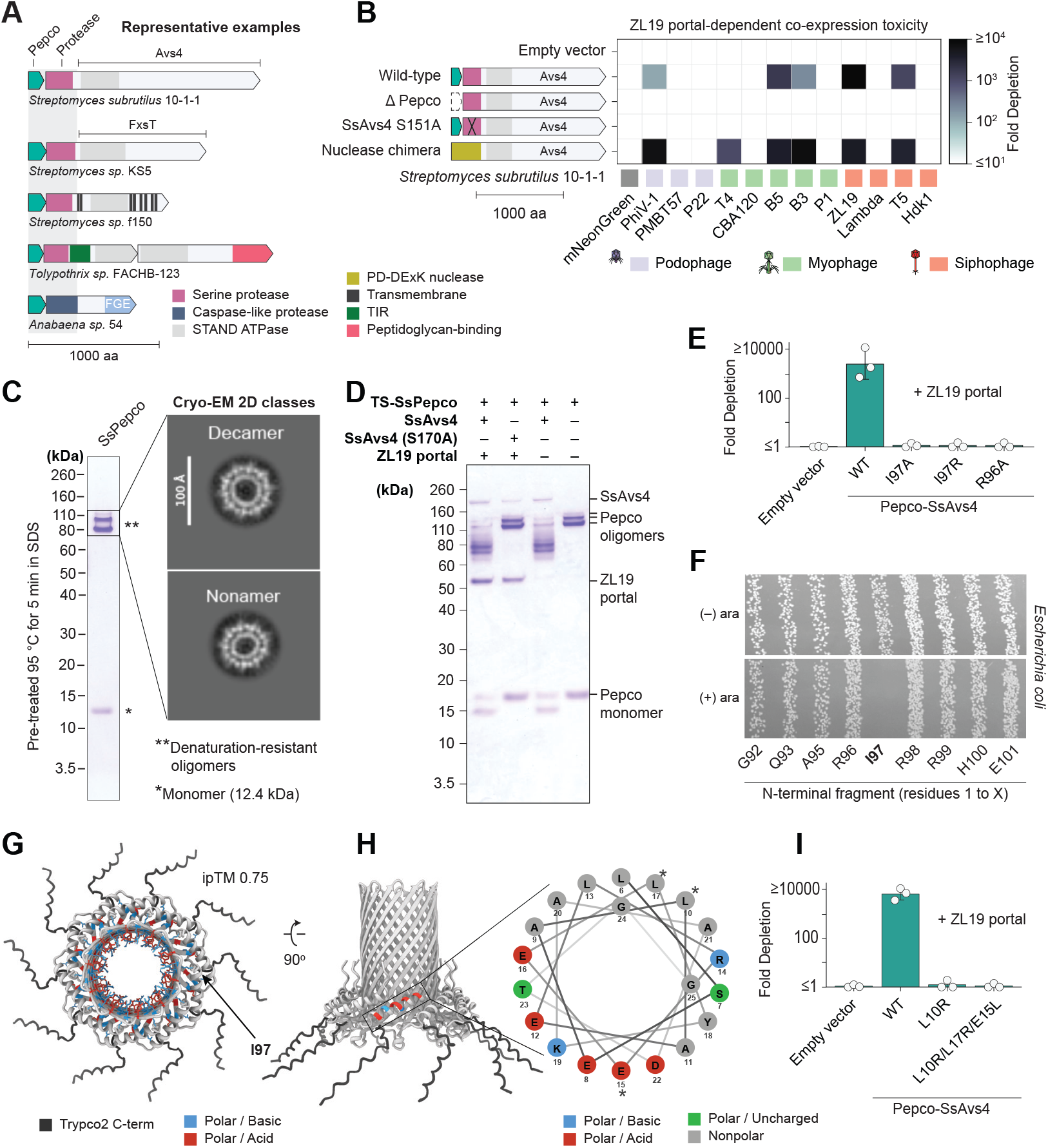
Pepco forms stable β-barrels that are proteolytically activated by C-terminal cleavage. (**A**) Representative genomic neighborhoods of Pepco–protease pairs. (**B**) *E. coli* co-expression toxicity of the *Streptomyces subrutilus* Pepco-protease (SsPepco–SsAvs4) defense system or its catalytic mutants with diverse phage portal proteins. The chimera is a synthetic fusion of the sensor domain from SsAvs4 and a nuclease-like effector domain from a homologous Avs4 in *Streptomyces exfoliatus*. (**C**) Purification of SsPepco, visualized by SDS-PAGE under denaturing conditions, shown with cryo-EM 2D classification of oligomeric fractions. (**D**) SDS-PAGE analysis of TwinStrep-tagged SsPepco cleaved by SsAvs4. (**E**) *E. coli* co-expression toxicity assay with ZL19 portal and the SsPepco–protease defense system, or mutations in the cleavage site residues I97 and R98 of SsPepco. (**F**) Expression toxicity assay of SsPepco fragments with shifted cleavage sites, with 0 indicating the cleavage position determined by mass spectrometry. (**G**) AlphaFold2 multimer model of a SsPepco decamer (ipTM 0.75). Basic and acidic residues on the β-strands are colored blue and red, respectively. The cleaved C-terminus of SsPepco is shown in dark gray. (**H**) Side view of the decameric SsPepco model, with a helical wheel diagram of the amphipathic helix at the base of the barrel. Stars indicate residues mutated to disrupt amphiphilicity. (**I**) Co-expression toxicity of SsPepco amphipathic residue mutations, in combination with SsAvs4 and the ZL19 portal.

SDS-PAGE analysis of recombinantly purified SsPepco revealed three high–molecular weight bands in addition to the expected 12.3 kDa monomer (Fig. 5C). Mass spectrometry confirmed that all bands corresponded to SsPepco, indicating the formation of unusually stable oligomers resistant to heat and detergent. Cryo-EM and structural modeling showed that these oligomers adopt symmetric β-barrel architectures (Fig. 5C). Two-dimensional cryo-EM class averages of top-down views revealed predominantly nonameric and decameric assemblies, with rare undecamers observed for affinity-tagged SsPepco (Fig. 5C, Supplementary Fig. 9A, and Supplementary Fig. 10A). These stoichiometries were consistent with the oligomers most confidently predicted by AlphaFold2 multimer (Supplementary Fig. 9B).

Recombinantly purified SsAvs4 cleaved both monomeric and oligomeric SsPepco into lower molecular weight fragments (Fig. 5D). Mass spectrometry revealed that a 2 kDa C-terminal tail of each SsPepco monomer is removed by SsAvs4 (Supplementary Fig. 11A–B and 9B). The Cterminal tail is rich in charged residues (40%) and proline (20%) and predicted to be unstructured (Supplementary Fig. 9B). Cleaved β-barrels remained predominantly nonameric and decameric (Supplementary Fig. 10B), suggesting that proteolysis remodels rather than disassembles the oligomer.

An isoleucine residue immediately upstream of the cleavage site is highly conserved among Pepco homologs associated with serine proteases. Substituting this residue (I97A or I97R) ablated the toxicity of SsPepco when co-expressed with SsAvs4 and the ZL19 phage portal (Fig. 5E). Conversely, expression of a truncated SsPepco ending precisely at Ile97 was sufficient to cause complete *E. coli* cell death under high arabinose induction (Fig. 5F). This effect was strictly position-dependent, as shifting the truncation site by even a single residue abolished toxicity. These results suggest that SsPepco activation requires specific proteolytic cleavage at Ile97.

Structural models revealed a strong enrichment of charged residues within the barrel lumen, while the barrel exterior is predominantly hydrophobic (Fig. 5G–H). Charged residues also cluster within the basal α-helices, which display an amphipathic organization with distinct charged and hydrophobic faces (Fig. 5H). These features are characteristic of membrane-active proteins. To test the role of the amphipathic helices, we introduced mutations that disrupt the amphipathic charge distribution. Scrambling the helix (L10R + L17R + E15A) or introducing a single substitution on the hydrophobic face (L10R) abolished toxicity (Fig. 5I). These results are consistent with SsPepco functioning as a protease-activated β-barrel pore that permeabilizes the membrane following Cterminal truncation.

### Cleaved EACC1 monomers oligomerize and disrupt lipid membranes

EACC1 is a small non-enzymatic protein encoded alongside cysteine proteases that are fused to diverse domains, including diversity-generating retroelements (DGRs)^31^ and Avs (Fig. 6A). Notably, EACC1–protease modules are fused to apparently non-defense components, such as DnaK-like chaperones and homologs of the type VII secretion protein EccC (Fig. 6A and Supplementary Fig. 12). We investigated a representative from *Ktedonobacter racemifer* DSM 44963, in which EACC1 is encoded upstream of a caspase-like protease domain fused to Avs12 (KrAvs12), which is activated by phage tail connector proteins. Co-expression of EACC1 and KrAvs12 with a phage tail connector induced strong cellular toxicity (Fig. 6B), and this effect was abolished by deletion of EACC1 or mutation of the protease catalytic cysteine (C152A).

**Figure 6.**
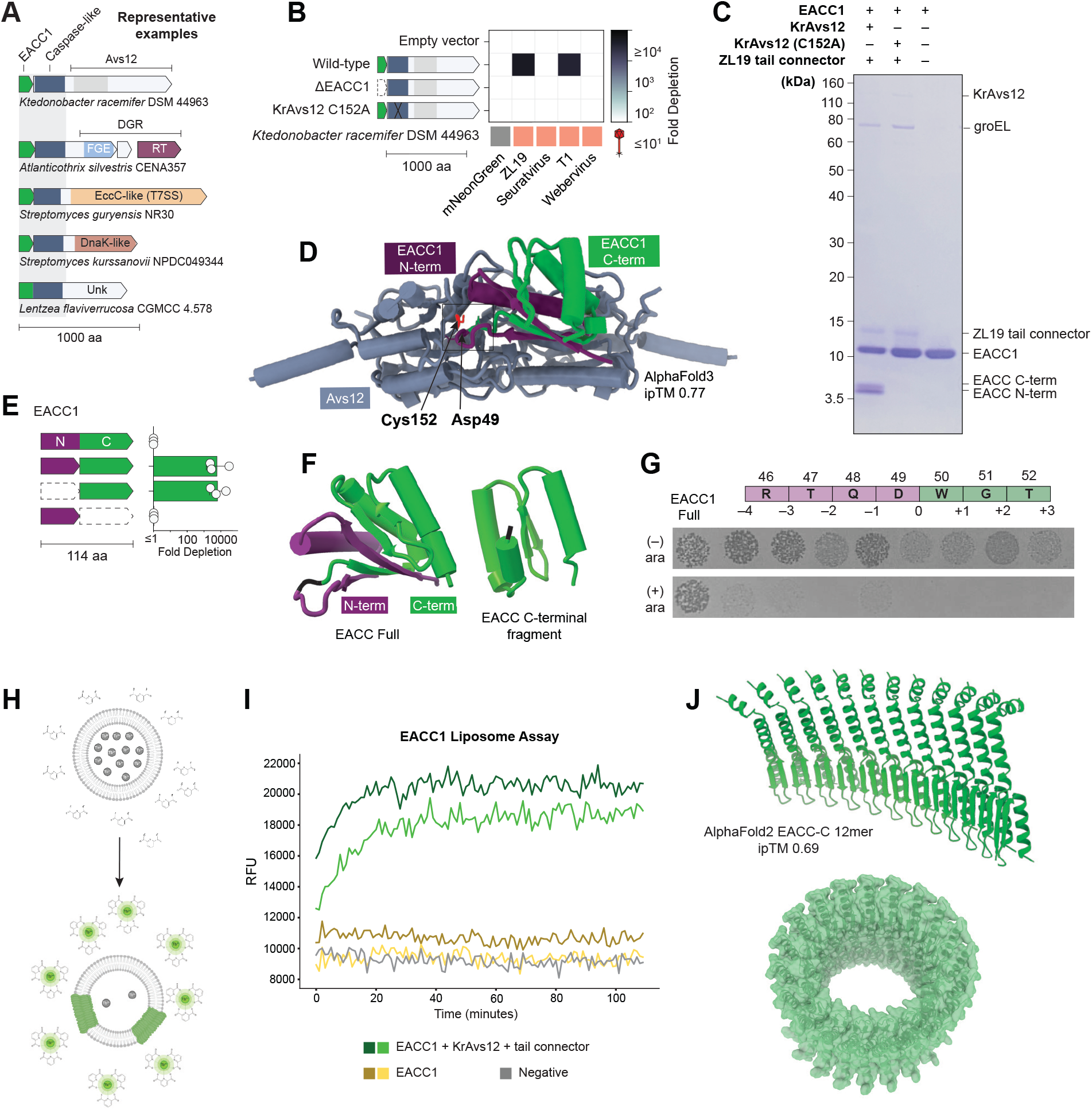
Cleaved EACC1 monomers oligomerize into a large membrane pore. (**A**) Representative genomic neighborhoods of EACC1–protease pairs. (**B**) *E. coli* co-expression toxicity of the *Ktedonobacter racemifer* EACC1-caspase (EACC1–KrAvs12) defense system or its catalytic mutants with diverse phage tail connector proteins. (**C**) SDS-PAGE of EACC1 cleaved by KrAvs12. (**D**) AlphaFold2 multimer model of EACC1 with a dimer of the KrAvs12 caspase domain (EACC1–caspase ipTM 0.77). (**E**) Expression toxicity assay of EACC1 fragment combinations in *E. coli* to mimic proteolytic cleavage. (**F**) AlphaFold3 model of a EACC1 monomer with the N-terminal domain in purple and the C-terminal domain in green. (**G**) Expression toxicity assay of EACC1 fragments with shifted cleavage sites, with 0 indicating the predicted cleavage position. (**H**) Schematic of a terbium-loaded liposome assay. (**I**) Time-course fluorescent measurement of liposomes with cleaved and uncleaved EACC1. (**J**) AlphaFold2 multimer model of C-terminal fragments of EACC1.

*In vitro* reconstitution with purified proteins revealed that KrAvs12 cleaves EACC1 into two fragments, a 7.1 kDa C-terminal fragment and a 5.7 kDa N-terminal fragment (Fig. 6C). Catalytic mutation of KrAvs12 (C152A) abolished cleavage (Fig. 6C). An AlphaFold3 model of EACC1 in complex with a KrAvs12 protease-domain dimer (ipTM 0.77) positioned the catalytic cysteine adjacent to a putative cleavage site at Asp49 (Fig. 6D). When expressed individually in *E. coli*, only the C-terminal fragment was toxic (Fig. 6E), consistent with its role as the downstream death effector. In contrast, the N-terminal fragment was not toxic, and structural modeling suggested that it forms a β sheet with two C-terminal β strands (Fig. 6F), supporting an autoinhibitory function. Shifting the cleavage site upstream or downstream preserved toxicity (Fig. 6G), suggesting EACC1 activation tolerates modest variation in cleavage position.

We next asked whether cleaved EACC1 might affect membrane permeability, as its autoinhibitory fragment is reminiscent of those of pore-forming proteins such as gasdermins^15,16,32^. To test this model, we performed a liposome leakage assay^15,33^ in which uncleaved or cleaved EACC1 was incubated with liposomes preloaded with terbium(III) ions that fluoresce upon release into external dipicolinic acid solution (Fig. 6H). Only EACC1 cleaved by KrAvs12 disrupted liposomes, consistent with a membrane permeabilization mechanism (Fig. 6I). Supporting this model, AlphaFold2 multimer predicted the formation of large oligomeric rings stabilized by β-strand interactions between adjacent protomers (Fig. 6J). Proteolytic cleavage may expose these β-strands, promoting oligomerization. Together, these findings support a model in which EACC1 assembles into a toxic membrane pore following proteolytic activation.

## Discussion

We investigated the mechanisms of four major prokaryotic protease–effector defense modules, each comprising a protease and its associated effector protein (Fig. 7). Biochemical, cellular, and structural analyses reveal that each effector encodes a zymogen or propeptide that is activated into a cell death toxin upon proteolytic cleavage. One effector is a DNase with an MBL-like fold that is autoinhibited by two linker regions; cleavage of the linkers by a phage-induced protease tetramer exposes the active site for Zn^2+^-dependent dsDNA degradation. These findings corroborate recent work on MBL by Tuck et al.^34^ A second effector is an α/β-fold hydrolase that requires double proteolytic cleavage to remove an inhibitory fragment and trigger cell death. Two additional effectors are small propeptides that form pore-forming toxins after cleavage; Pepco is a pre-assembled β-barrel that becomes toxic once processed, while EACC1 is an autoinhibited monomer that assembles into a membrane pore only after proteolytic cleavage.

**Figure 7.**
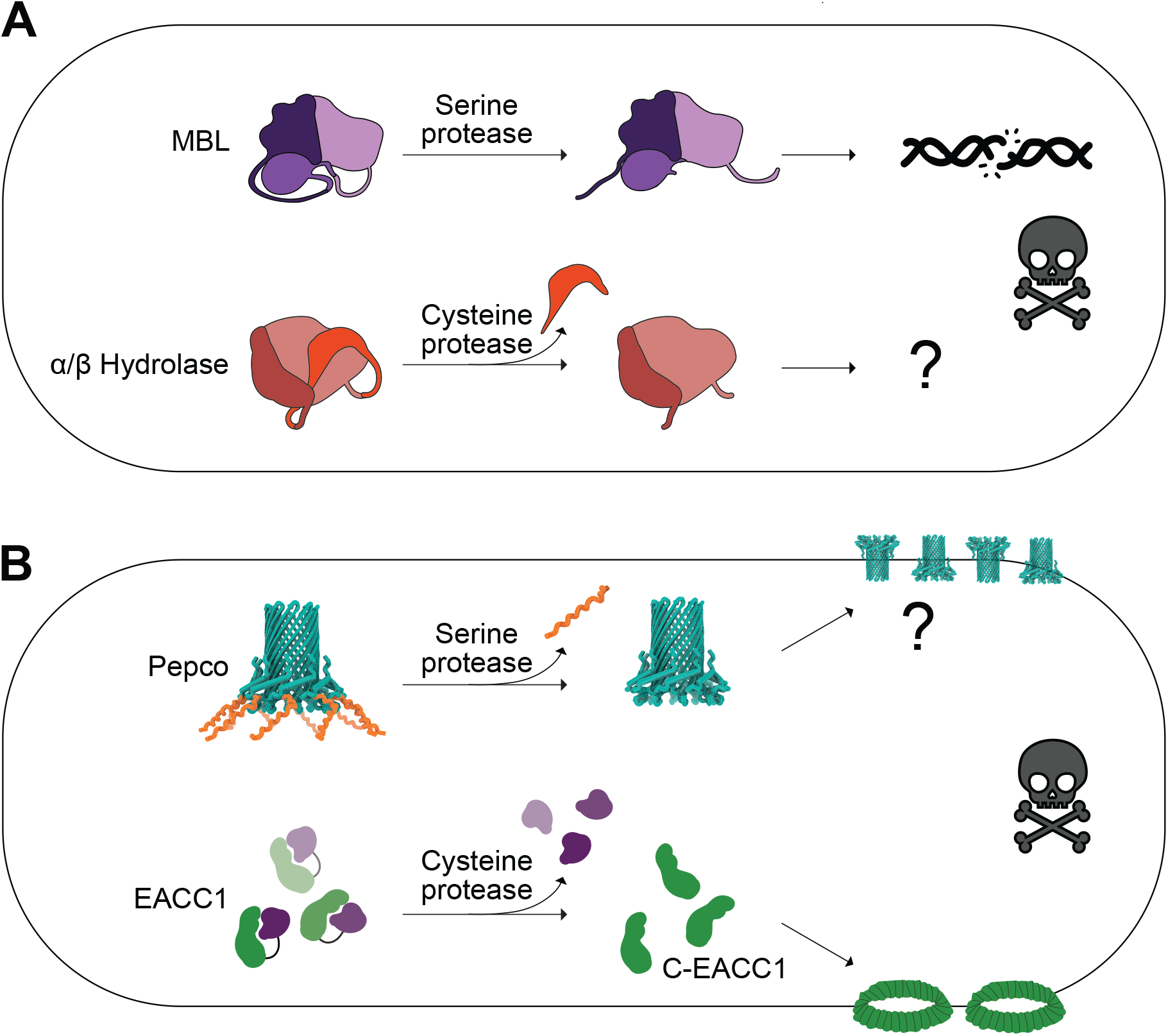
Proteolytic activation of diverse effectors is a conserved immune strategy. Graphical summary of (**A**) two zymogens and (**B**) two membrane pores activated by associated proteases.

Mining prokaryotic genomes revealed that protease–effector modules are widely distributed, with tens of thousands of homologs detectable in the NCBI non-redundant protein database. Conserved pairing of effectors with protease suggests an evolved dependency on proteolytic cleavage and conservation in the cell death mechanisms for each effector family. Moreover, each protease–effector module has recombined with a wide array of immune sensors, enabling defense against diverse phage infection signals. Remarkably, several EACC1–protease homologs are fused to a DnaK-like chaperone that shares 50% sequence identity with canonical DnaK, a prokaryotic Hsp70 chaperone that assists protein folding and protects cells from thermal or oxidative stress^35,36^. The association of DnaK-like domains with EACC1 modules raises the possibility that protease–effector activation can function in cellular stress responses beyond phage defense. Interestingly, the DnaK-like domain lacks the C-terminal lid helices that normally trap unfolded substrates^37,38^, suggesting it may sense misfolded proteins without refolding (Supplementary Fig. 12). Alternatively, or additionally, the DnaK-like domain may have evolved to respond to stresses associated with phage infection. Together, these findings suggest that protease–effector modules are evolutionarily conserved and functionally adaptable, capable of integrating multiple signals to regulate targeted cell death.

Proteases are also central regulators in the innate immune systems of eukaryotes, triggering cascades of inflammation, apoptosis, and pyroptosis in response to an activation signal^39,40^. For instance, caspases control cytokine release, pore formation through gasdermins, and DNA degradation through caspase-activated DNase (CAD)^18,40–42^. Our findings support the conclusion that a similar regulatory logic is widespread in prokaryotes. For example, EACC1 monomers oligomerize into a pore only after proteolytic cleavage, similar to the activation mechanism seen in mammalian perforin and gasdermin pores^18,32,42^. Likewise, although activation of Pepco through C-terminal cleavage of pre-formed β-barrels represents a distinct mechanism of pore formation, the Pepco barrels resemble miniature versions of the mammalian membrane attack complex/perforin (MACPF) family^18^. Moreover, we find that protease-dependent regulation of nuclease activity is another conserved immune mechanism. In mammals, for example, caspase-3 cleaves the inhibitor of CAD, allowing CAD to dimerize and degrade nuclear dsDNA^43^. Such regulation is paralleled in MBL, which represents a novel example of a nuclease in prokaryotic immunity that requires proteolytic activation. MBL is also the first zymogen known to require two distinct cleavage events, with all resulting fragments required for its active form.

For SsPepco and EACC1, protease activity appeared constitutive *in vitro* despite showing trigger-dependent toxicity in cells. This discrepancy might reflect differences in protein concentration, molecular crowding, or other intracellular factors not recapitulated in purified systems; indeed, constitutive *in vitro* activity has also been observed for other protease-based bacterial defense systems, which nonetheless accurately reproduce cleavage specificity^15,33^. The close agreement among our site-specific *in vitro* cleavage patterns, cellular toxicity assays, and high-confidence structural models strongly supports accurate identification of protease function and substrate cleavage sites. Additionally, cryo-EM micrographs of SsPepco were limited to top-down views despite extensive variation in sample preparation conditions, preventing experimental reconstruction of a high-resolution threedimensional structure. Nevertheless, this pronounced orientation bias enabled clear classification of distinct oligomeric states.

Beyond the four protease–effector modules we have described here, we also investigated an uncharacterized caspase-like protease with no associated effector. A homolog found in *E. coli* 27-18K conferred resistance to phages PhiV1, T4, and T2 when heterologously expressed in *E. coli* K-12 (Supplementary Fig. 13), and this protection was abolished upon mutation of the catalytic cysteine (C131A). Effector-independent proteases may confer defense through broad proteolysis of the host proteome, or by specific cleavage of a conserved host factor or phage protein (Supplementary Fig. 13D). Further work will be required to determine how these caspases lacking dedicated effector partners mediate phage defense.

Collectively, our findings expand the repertoire of mechanisms by which bacteria resist infection, highlighting protease–effector modules emerge as a versatile and conserved strategy for immune regulation. The shared logic between these bacterial systems and eukaryotic immunity highlights a unifying principle in which toxic proteins are maintained in an autoinhibited state until activated by proteolytic cleavage, which may enable tight regulation and precise signal amplification in response to a wide range of cellular threats.

## Materials and Methods

### Bioinformatics analyses

Serine protease-associated MBL homologs (Fig. 1C and Supplementary Fig. 1) were identified via PSI-BLAST^44^ searches against NCBI nr in November 2024. After several rounds of manual curation to remove truncated sequences and hits outside of the serine protease-associated clade, a final set of 2,689 verified MBL sequences was obtained. For each MBL, the core sequence was concatenated with its associated protease sequence, except for 15 non-protease-associated MBLs, which were used with the MBL core sequence alone. The concatenated sequences were clustered at 90% identity and 95% coverage using MMseqs2 (ref.^45^) (–min-seq-id 0.9 -c 0.95 –cov-mode 0), resulting in 1,502 clusters. One representative from each cluster was selected for further analysis. A multiple sequence alignment (MSA) of the representative MBL sequences (excluding the protease sequence) was generated using MAFFT v7.520 (ref.^46^) with global pairwise alignment (–maxiterate 1000 –globalpair), then trimmed with trimAl 1.2 (ref.^47^) using a gap threshold of 0.25 (-gt 0.25). Phylogenetic trees were constructed from the trimmed MSAs using IQ-TREE 1.6.12 (ref.^48^) with the LG+G4 model and 2000 ultrafast bootstrap replicates (parameters -nstop 500 -safe -nt 4 -bb 2000 -m LG+G4). Sensor domains of the associated proteases were classified through sequence clustering, guided by HHpred^30^ analyses and AlphaFold3 (ref.^49^) structural predictions, and taxonomic classifications were retrieved from the NCBI taxonomy database. The tree was visualized using iTOL v6 (ref.^50^). A similar procedure was applied for the serine protease tree (Supplementary Fig. 1A). A tanglegram was visualized with ladderized phylogenetic trees of MBL and serine protease using the Baltic Python library (Dudas, GitHub). Tip nodes were colored by hierarchically ordered sequence clusters from MMseq2 at 30% identity and 70% coverage for MBL and 20% identity and 50% coverage for protease (Supplementary Fig. 1B).

### Cloning

Genes were chemically synthesized (Twist Bioscience or Integrated DNA Technologies) or amplified by PCR from phage genomic DNA. Plasmids were constructed with Gibson assembly and transformed into *E. coli* Stbl3 or Mach1 (ThermoFisher), and colonies were cultured in Terrific Broth at 37 °C with carbenicillin (100 µg/mL) or chloramphenicol (25 µg/mL). Plasmids were purified using QIAGEN miniprep buffers and 96-well DNA & RNA binding plates (Epoch Life Science) and were fully sequence verified by Tn5 tagmentation and deep sequencing, as described previously^4,51^. Phage genes were cloned into barcoded pBAD expression vectors with ampicillin resistance. Defense genes were cloned into a pACYC184-like vector backbone with chloramphenicol resistance. For genes from non-Enterobacteriaceae sources, coding sequences were codon-optimized for expression in *E. coli* and placed downstream of a pLac promoter. For the novel *E. coli*–derived caspase system (Supplementary Fig. 13), the native promoter and coding sequence were retained.

### Plaque assays

*E. coli* ATCC25404 harboring a defense system, a mutant, or an empty vector was grown in Terrific Broth to an optical density at 600 nm (OD_600_) of 0.8 to 1.0. A 1-mL aliquot was mixed with 49 mL of top agar (7 g/L agar, 15.5 g/L LB Lennox, 4.5 g/L NaCl) supplemented with 25 µg/mL chloramphenicol, poured into a 15 cm Petri dish, and allowed to solidify at room temperature. Ten-fold serial dilutions of phage in deionized water were spotted on top, with 3 µL per spot. Plates were incubated for 17 h at 37 °C, then imaged in the dark over a white backlight using a Nikon D7500 DSLR camera. Images were rotated, scaled, cropped, and quantified using custom Python and ImageJ^52^ scripts.

### Co-expression toxicity screens

Barcoded pBAD vectors encoding phage genes were pooled to generate a library with balanced representation. Control plasmids expressing mNeonGreen were spiked in as normalization controls. The library was transformed into *E. coli* NovaBlue(DE3) cells expressing protease–effector modules or their corresponding mutants. After 1 h outgrowth at 37 °C in SOC, cells were plated on LB agar containing 100 µg/mL carbenicillin, 25 µg/mL chloramphenicol, and arabinose (0.05% for VcMBL, AtMBL, and EACC1; 2% for SsPepco). Plates were incubated for 12 h at 37 °C, and plasmids from surviving colonies were isolated by miniprep. Barcode regions were amplified by PCR and sequenced on an Illumina MiSeq. Read counts were normalized to the number of mNeonGreen barcode reads per sample, and fold depletion for each gene was calculated by comparing barcode abundance in the treatment to that in the empty vector control.

### Expression toxicity screens

Barcoded pBAD vectors encoding protease-associated effector fragments were cloned and pooled to generate a library with balanced representation, as described previously^5^. To model proteolytic activation of zymogens requiring two cleavages, the N-, middle (M), and C-terminal fragments of MBL and LcHydrolase were expressed individually, in pairs, or all three together. Singlesite cleavage was modeled by expressing N with M+C or C with N+M, whereas expression of the full-length protein represented the uncleaved state. For proteins requiring a single proteolytic cut (EACC1 and Pepco), N- and C-terminal fragments were expressed individually or together, with the fulllength protein again representing the uncleaved state. To test cleavage specificity at the scissile bond, alternative fragment sets were generated by shifting the cleavage junction 3 to +3 residues relative to the native site. Control plasmids expressing mNeonGreen were spiked in for normalization.

The pooled library was transformed into *E. coli* NovaBlue(DE3). After 1 hr outgrowth at 37 °C in SOC, cells were plated on LB agar containing 100 µg/mL carbenicillin with and without arabinose (0.05% for VcMBL, AtMBL, and EACC1 fragments; 2% for SsPepco and LcHydrolase fragments). Plates were incubated for 12 h at 37 °C, and plasmids from surviving colonies were isolated by miniprep. Barcode regions were PCR-amplified and sequenced on an Illumina MiSeq. Read counts were normalized to the number of mNeonGreen barcode reads per sample, and fold depletion for each gene was calculated by comparing barcode abundance under arabinose conditions to that under no-arabinose conditions.

### Protein purification

Plasmids encoding MBL-TwinStrep (TS), VcAvs2-TS, TS-Pepco, TS-SUMO-Pepco, SsAvs4His_6_, TS-SUMO-EACC, TS-SUMO-KrAvs12, TS-SUMO-ZL19 tail connector, His_6_-LcHydrolase, TS-SUMO-LcHydrolase, and TS-SUMO-LcCaspase were transformed into *E. coli* Rosetta(DE3) (Fisher Scientific). The T4 terminase-TS plasmid was transformed into *E. coli* BL21(DE3) (New England Biolabs) containing pLysS. Transformations were plated on LB agar containing 100 µg/mL carbenicillin and 25 µg/mL chloramphenicol. After overnight growth at 37 °C, single colonies were inoculated into Terrific Broth (TB) with the appropriate antibiotics and cultured overnight at 37 °C. Starter cultures were diluted 1:200 into fresh, identically supplemented TB and grown at 37 °C to an OD_600_ of ∼0.5. Protein expression was induced with 0.1 mM isopropyl-β-D-1-thiogalactopyranoside (IPTG), and growth temperature was lowered to 18 °C. All cultures were subsequently grown for 16 h, except TS-SUMO-KrAvs12, which was grown for 6 h to reduce chaperone contamination. Cells were pelleted by centrifugation and stored at 80 °C until use.

Pellets were resuspended in 40 mL lysis buffer (250 mM NaCl, 50 mM Tris-HCl pH 7.5, 5% glycerol, 5 mM β-mercaptoethanol) and lysed at 28,000 PSI in an LM20 Microfluidizer (Microfluidics). Lysates were clarified by centrifugation at 39,000 x g for 45 min at 4 °C, and the supernatant was incubated with Strep-Tactin Superflow Plus resin (QIAGEN) for 1 h at 4 °C. Resin-bound proteins were washed six times with lysis buffer on a Poly-Prep chromatography column (Bio-Rad). Elution and tag processing were as follows: TS-tagged proteins were eluted with lysis buffer containing 5 mM desthiobiotin; His-tagged proteins were eluted stepwise with lysis buffer containing 20, 50, and 500 mM imidazole; TS–SUMO fusions were incubated overnight at 4 °C in lysis buffer with bdSENP1 protease. Proteins exhibiting contaminant bands were further purified by size-exclusion chromatography on a Superose 6 Increase 10/300 GL or Superdex 200 column (Cytiva) equilibrated in lysis buffer using an ÄKTA Explorer (Cytiva).

### In vitro protease assays

Effector proteins were incubated with approximately 0.1 molar equivalents of their corresponding protease and, when applicable, the corresponding phage protein trigger in 20 µL reactions at 37 °C. Specifically, VcMBL (5 µM) was incubated for 5 min with or without 500 nM VcAvs2 and 500 nM T4 terminase in the presence of 1 mM ATP, 5 mM Mg^2+^, and 100 µM Zn^2+^. SsPepco (10 µM) was incubated for 30 min with or without 1 µM SsAvs4 and 3 µM ZL19 portal in the presence of 1 mM ATP and 5 mM Mg^2+^. EACC1 (25 µM) was incubated for 30 min with or without 500 nM KrAvs12 and 4 µM ZL19 tail connector, also in the presence of 1 mM ATP and 5 mM Mg^2+^. Lastly, LcHydrolase (10 µM) was incubated with or without 1 µM LcCaspase in the presence of 1 mM ATP across a time course of 0, 5, 10, 30, 60, and 240 min. All reactions were terminated by addition of Laemmli buffer and boiled at 95 °C for 5 min. Proteins were resolved on 4-12% NuPAGE Bis-Tris gels (Thermo Fisher Scientific) and visualized by Coomassie staining using an eStain L1 system (GenScript).

### In vitro nuclease assay

VcMBL (5 µM) was incubated with 500 nM VcAvs2 and 500 nM T4 terminase in the presence of 1 mM ATP and 5 mM Mg^2+^ at 37 °C for 5 min. A portion of this cleavage reaction was then added to 150 ng of doublestranded DNA (dsDNA) in buffer containing 100 µM Zn^2+^, bringing the final VcMBL concentration to ∼100 nM. After a 5-min incubation at 37 °C, the mixture was purified using a PCR cleanup kit (QIAGEN) and analyzed by electrophoresis on an E-Gel EX 2% agarose gel (Thermo Fisher Scientific). The remaining protease reaction was quenched with Laemmli buffer, boiled at 95 °C for 5 min, and analyzed by SDS-PAGE on a 4-12% Bis-Tris gel (Thermo Fisher Scientific), followed by Coomassie staining.

### Liposome permeability assay

A total of 15 mg 1,2-dioleoylsn-glycero-3-phosphocholine (DOPC) in 300 µL chloroform was dried in a round-bottom glass tube by rotation under argon gas. The dried lipids were resuspended in pentane, redried under argon with rotation, and stored overnight in a vacuum desiccator. Lipids were then resuspended in 800 µL terbium resuspension buffer (15 mM TbCl_3_, 20 mM HEPES-KOH pH 7.5, 50 mM sodium citrate, 150 mM NaCl)^15,33^. Crude liposomes were prepared by 10 cycles of vortexing (1 min), freezing in liquid nitrogen, and thawing in a room-temperature water bath. To improve uniformity, the suspension was pulse centrifuged and subjected to 15 additional cycles of liquid sonication (1 min) followed by resting in a water bath (1 min). Final liposomes were stored at 4 °C in 14 kDa MWCO dialysis tubing (Thermo Fisher Scientific) in dialysis buffer (20 mM HEPES-KOH pH 7.5, 50 mM sodium citrate, 150 mM NaCl) for 4 days.

Proteins were buffer-exchanged into dialysis buffer supplemented with 5 mM β-mercaptoethanol (BME). EACC1 (10 µM) was incubated with or without KrAvs12 (1 µM) protease and ZL19 tail connector (1 µM) in the presence of 1 mM ATP and 5 mM Mg^2+^ at 37 °C for 30 min. Replicates of this cleavage reaction were performed in parallel using independently purified preparations of EACC1, KrAvs12, and ZL19 tail connector.

Cleavage reactions were then supplemented with 1 µL of 100x dipicolinic acid (2 mM DPA). Each 70 µL cleavage reaction was added to 30 µL of DOPC liposomes, mixed by two cycles of pipetting, and immediately transferred to a Tecan Infinite M Nano plate reader. Fluorescence of the Tb^3+^-DPA complex was monitored for 1 h with measurements taken every minute by excitation and emission at 276 nm and 545 nm, respectively^15,33^.

### Cryo-EM

A total of 3 µL of SsPepco (5 mg/mL) was applied to freshly glow-discharged Au R1.2/1.3 300 mesh Quantifoil grids and vitrified in liquid ethane using a Vitrobot Mark IV (Thermo Fisher Scientific) operated at 4 °C and 100% humidity. Blotting parameters were as follows: 5 s wait time, 3 s blot time, and blot force 3. Grids were clipped and loaded into a 200 kV Thermo Fisher Glacios Cryo-TEM equipped with a K3 Summit direct electron detector (Gatan) operating in counting mode at a physical pixel size of 0.92 Å. Images were recorded using SerialEM, with each movie consisting of 50 frames collected over a total exposure time of 2.5 s, a total electron dose of 50 electrons per Å2, and a defocus range of 1.2 to 2.0 µm. A total of 525 movies were processed in cryoSPARC v4.4 (ref.^53^) for motion correction, patch CTF estimation, template-based particle picking, and particle extraction using a 64-pixel box size and 4×4 binning. Initial extraction yielded 1,031,283 particles. After three rounds of 2D classification, 482,509 particles with well-defined structural features were retained.

## Supporting information

MSAs and trees related to MBL

## ACKNOWLEDGEMENTS

We thank Fanny Sunden and Siyuan Du for valuable discussion; Collin Chiu for assistance with plaque assays; Jim Zhang for assistance with liposome preparation; Bharti Singal at Stanford cEMc for assistance with EM data collection; Ross Tomaino for mass spectrometry analysis (Taplin Biological Mass Spectrometry Facility, Harvard Medical School); Joel Nott for Edman degradation (Protein Facility, Iowa State University Office of Biotechnology); Owen Dunkley and Sayeh Gorjifard for manuscript feedback; and the entire Gao lab for support and advice. S.A.E. is supported by an NIH Genetics and Developmental Biology Training Grant (NIGMS, grant number 5T32GM141828) and the Bio-X Bowes Fellowship. L.F. is supported by the G. Harold and Leila Y. Mathers Foundation and NIH R35GM153424. Y.L. is supported by the NSF Graduate Research Fellowship Program DGE-2146755. Y.T.L. is supported by Knight-Hennessey Scholars at Stanford University. D.H. is supported by NSF MCB2322069. A.G. is supported by the G. Harold and Leila Y. Mathers Foundation, the Esther Ehrman Lazard Faculty Scholars Program, and Stanford Bio-X.

## AUTHOR CONTRIBUTIONS

S.A.E. and A.G. designed experiments. S.A.E. carried out the genetic screens and phage plaque assays. S.A.E., Y.L., and K.W. performed molecular cloning and *in vitro* biochemistry. J.Z. and S.A.E. performed cryo-EM. A.G., S.A.E., and Y.T.L. performed phylogenetic analyses. A.G. conceived and supervised the research. S.A.E. and A.G. wrote the manuscript with input from all authors.

