## Supplementary Figures for "Proteolytic activation of diverse antiviral defense modules in prokaryotes"

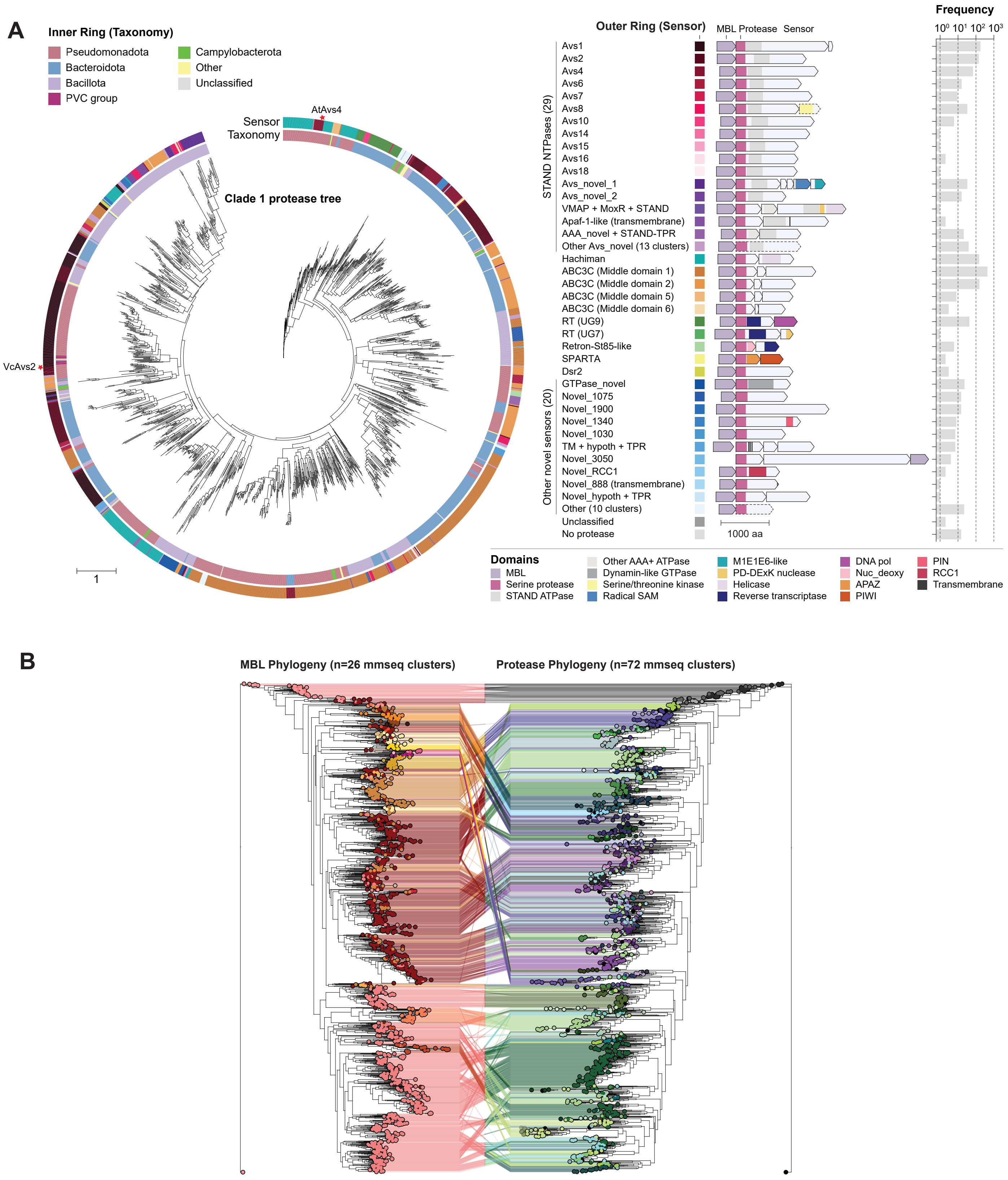


**Supplementary Figure 1. Phylogeny of clade 1 MBL-associated proteases (related to Figure 1).** (**A**) Maximum likelihood phylogenetic tree of the trypsin-like protease domain associated with clade 1 MBLs. (**B**) Tanglegram displaying flattened MBL (left) and protease (right) phylogenies inferred by maximum likelihood analysis of multiple sequence alignments, with tip colors representing hierarchically ordered sequence clusters obtained with MMseq2.

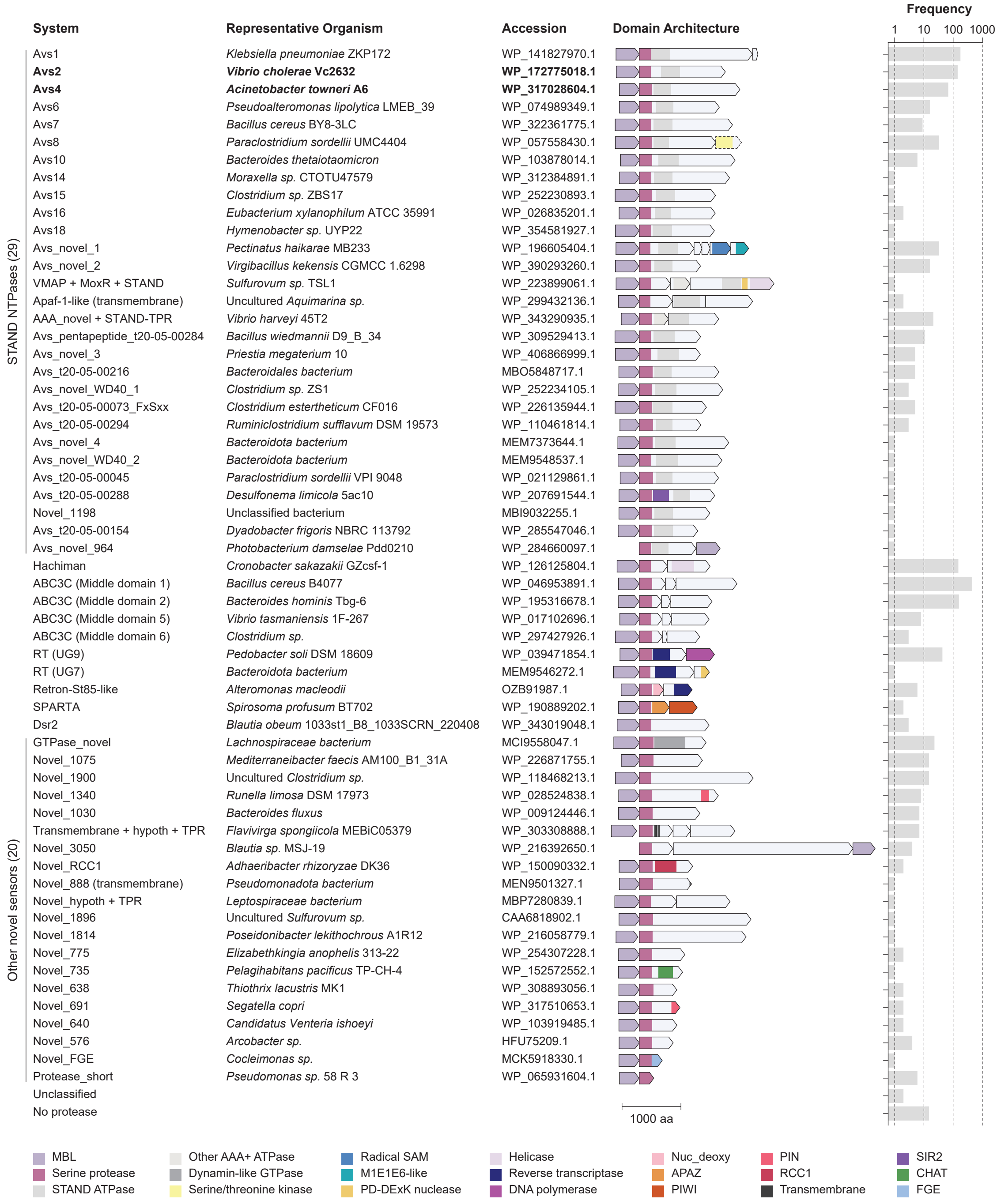


**Supplementary Figure 2. Representative loci of MBL homologs associated with trypsin-like serine proteases (related to Figure 1).** Representative loci of MBL homologs associated with trypsin-like serine proteases.

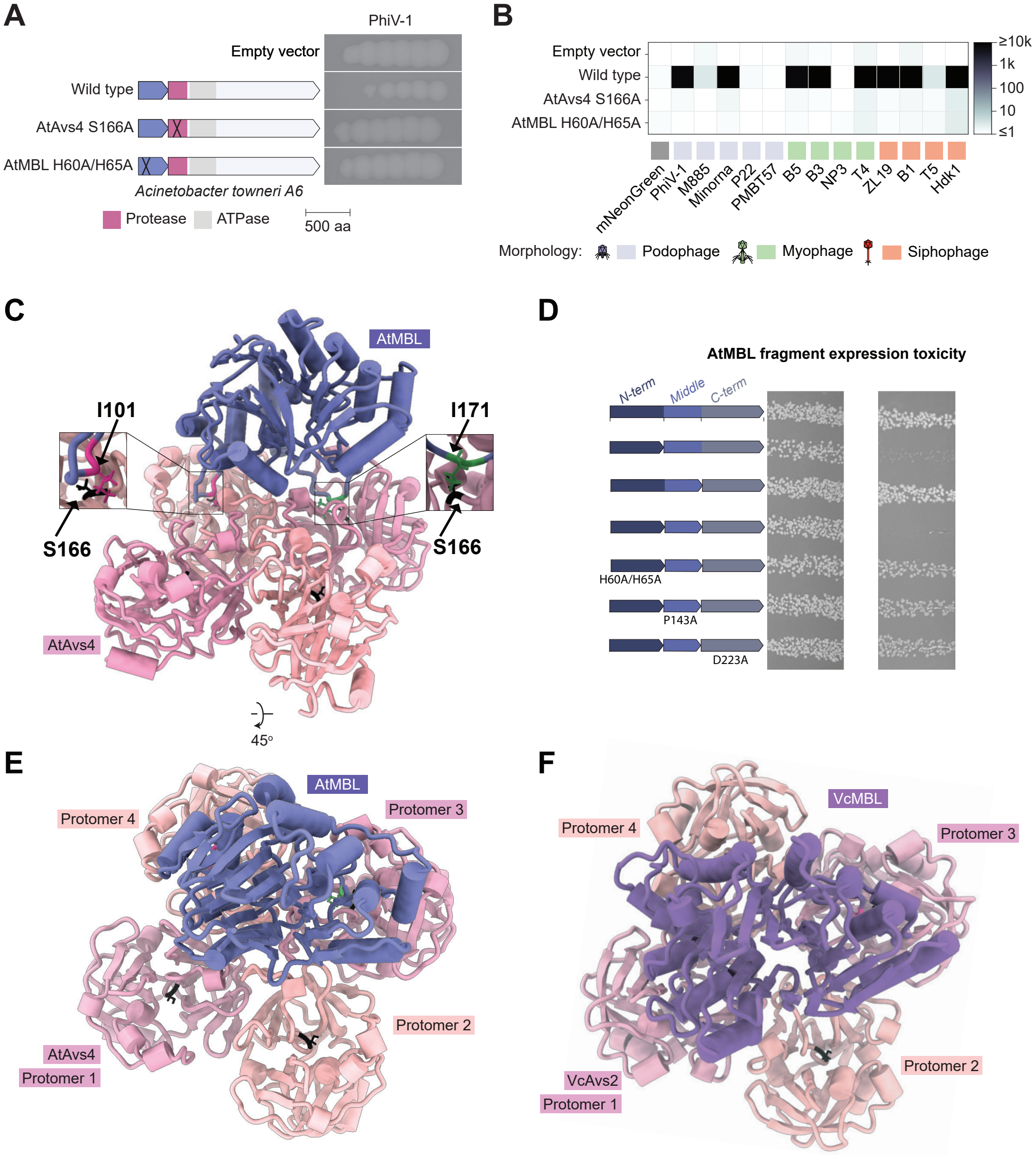


**Supplementary Figure 3. Proteolytic cleavage of an Avs4-associated MBL from *Acinetobacter towneri* (AtMBL) (related to Figure 2 and 3).** (**A**) Heterologous reconstitution of *Acinetobacter towneri (*AtAvs4) and its associated MBL (AtMBL) in *E. coli*. Plaque assay spots show 10-fold serial dilutions of phage PhiV-1 on strains expressing wild-type or catalytic mutants of AtMBL (H60A/H65A) and AtAvs4 protease (S166A). (**B**) *E. coli* co-expression toxicity of the AtMBL-protease defense system or its catalytic mutants with diverse phage portal proteins. (**C**) AlphaFold2 multimer model of AtMBL in complex with a C4-symmetric AtAvs4 protease tetramer, with catalytic residues positioned at predicted cleavage sites on AtMBL. (**D**) Co-expression toxicity assay of AtMBL fragment combinations in *E. coli* to mimic proteolytic cleavage. Protease interactions with AtMBL and VcMBL, showing (**E**) AtMBL bound to two adjacent protease protomers and (**F**) VcMBL bound to two diagonally opposed protomers.

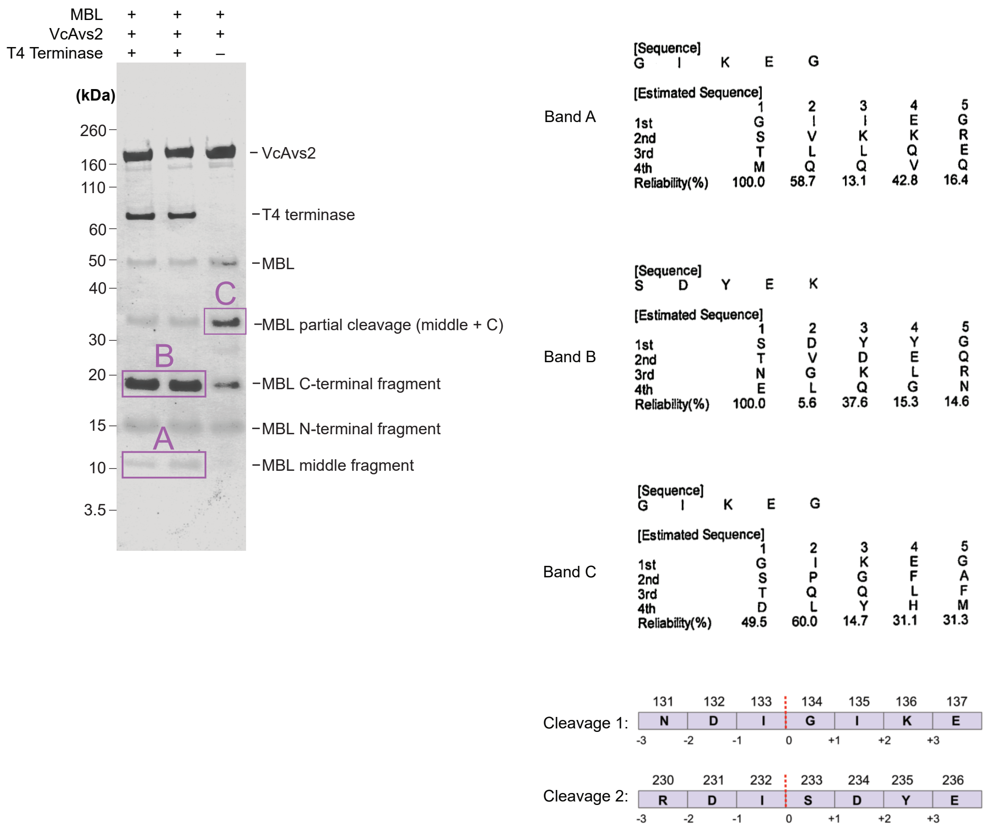


**Supplementary Figure 4. Edman degradation of VcMBL fragments after proteolytic cleavage (related to Figure 2).**

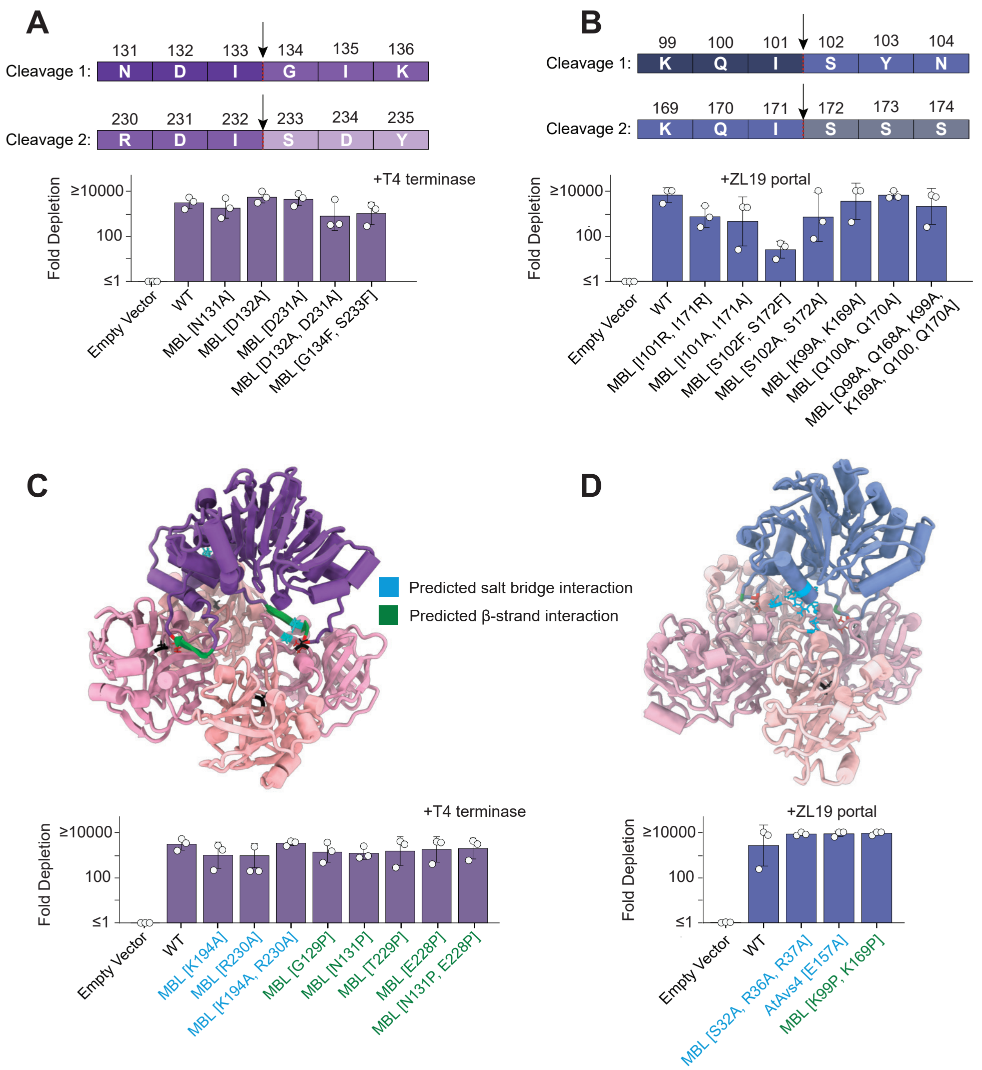


**Supplementary Figure 5. Mutational analysis of the Avs–MBL interface (related to Figure 2).** Mutations of the conserved cleavage motif in (**A**) VcMBL and (**B**) AtMBL. Alanine substitutions targeting salt bridges (blue) and β-strands (green) at the interface between (**C**) VcMBL and its associated protease, and (**D**) AtMBL and its protease.

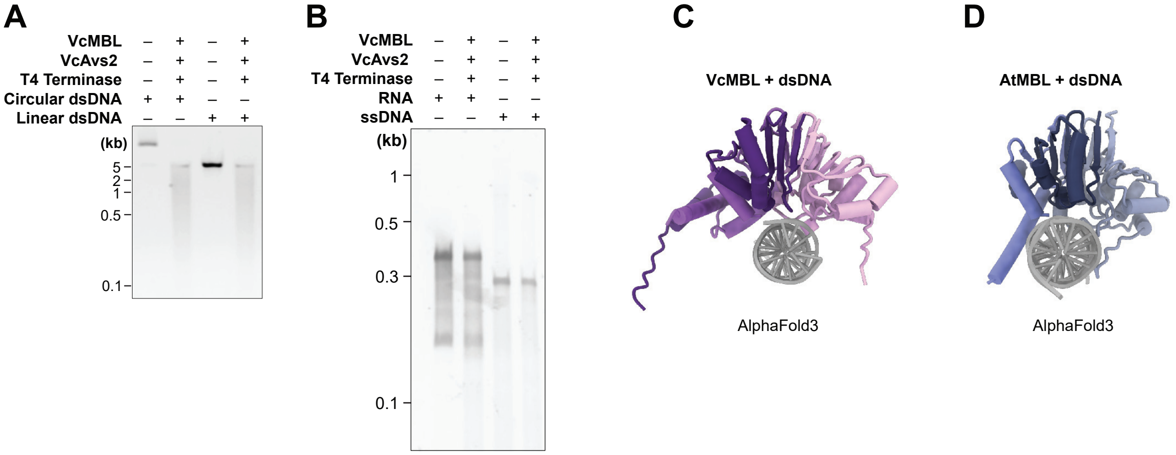


**Supplementary Figure 6. VcMBL is a double-stranded DNA endonuclease (related to Figure 3).** (**A**) Agarose gel of linear and circular dsDNA substrates incubated with cleaved VcMBL. (**B**) Agarose gel of RNA and ssDNA substrates incubated with cleaved VcMBL. (**C**) AlphaFold3 model of VcMBL in complex with dsDNA. (**D**) AlphaFold3 model of AtMBL in complex with dsDNA.

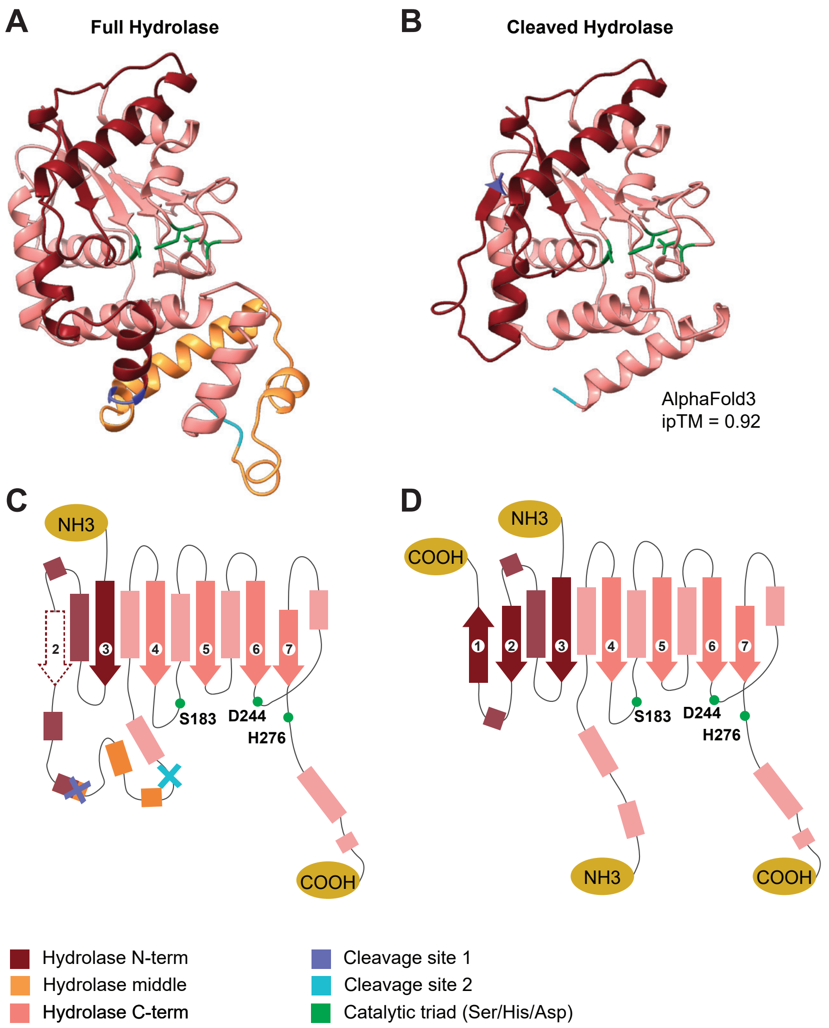


**Supplementary Figure 7. Comparison of the structures of full and cleaved LcHydrolase (related to Fig 4).** AlphaFold3 models of (**A**) wild-type LcHydrolase in the pre-cleavage state (full) and (**B**) N-terminal and C-terminal fragments of cleaved LcHydrolase. Secondary structure diagram of (**C**) full LcHydrolase and (**D**) cleaved LcHydrolase.

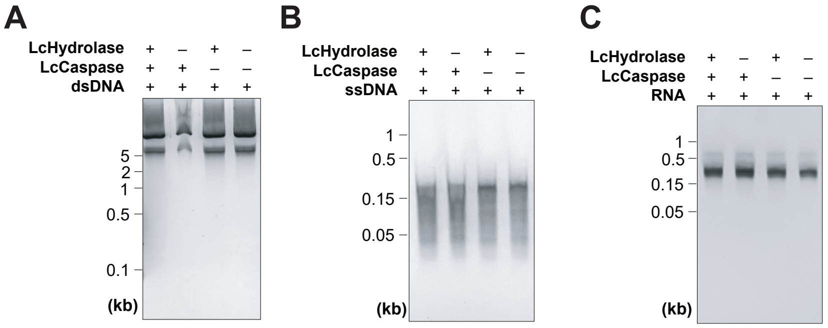


**Supplementary Figure 8. Activity of LcHydrolase on nucleic acid substrates (related to Figure 4).** Agarose gels showing incubation of cleaved LcHydrolase with (**A**) double-stranded DNA, (**B**) single-stranded DNA, and (**C**) RNA.

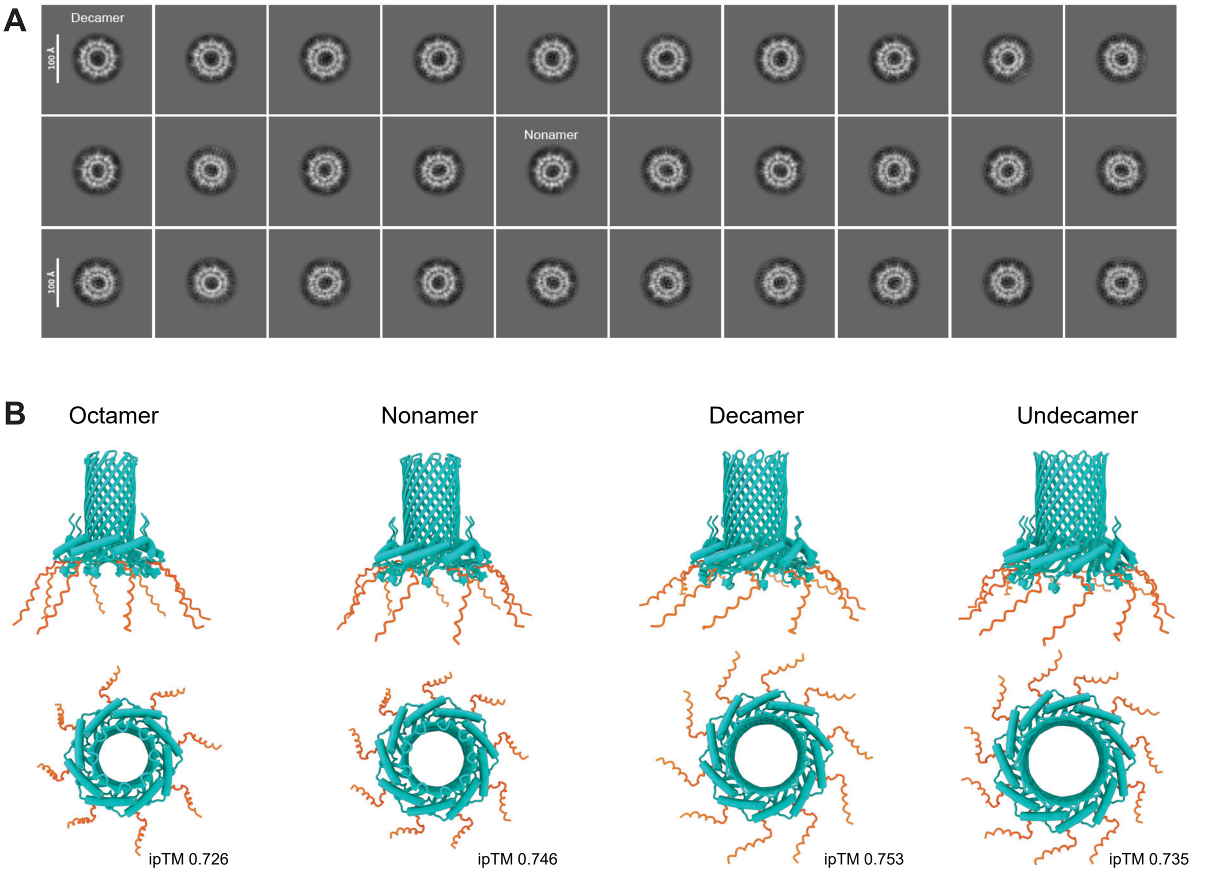


**Supplementary Figure 9. Pepco is predominately a nonamer and decamer (related to Figure 5).**
(**A**) Cryo-EM 2D class averages of top views of the Trypco2 β-barrel. (**B**) AlphaFold2 multimer models of 8- to 11-mer SsPepco oligomers. Orange C-termini indicate the fragment cleaved by SsAvs4.

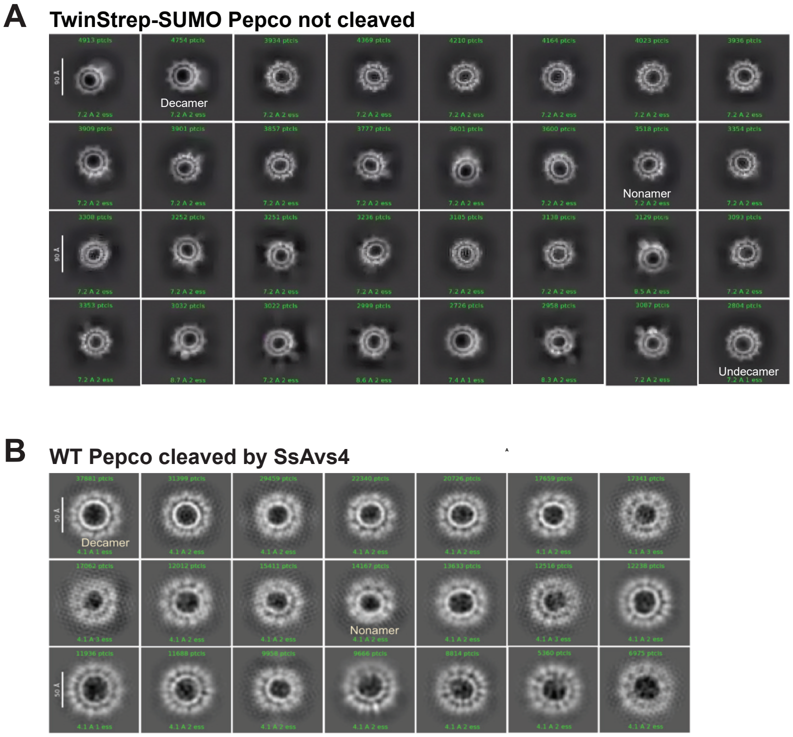


**Supplementary Figure 10. The Pepco β-barrel persists after N-terminal tagging or protease cleavage (related to Figure 5).** Cryo-EM 2D class averages of (**A**) uncleaved Trypco2 β-barrels with an N-terminal TwinStrep-SUMO tag on each monomer and (**B**) Trypco2 β-barrels following cleavage by SsAvs4.

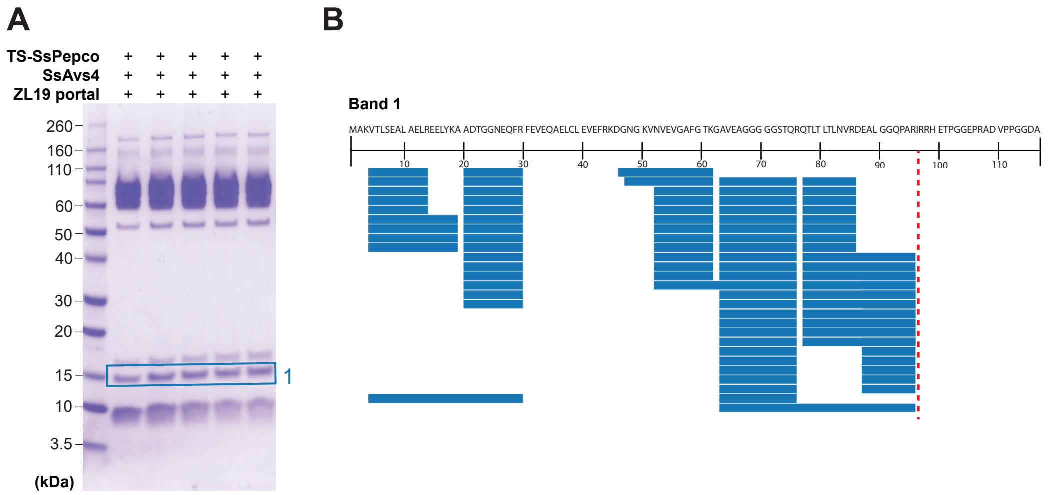


**Supplementary Figure 11. Mass spectrometry analysis of SsPepco cleavage products (related to Figure 5).** (**A**) SDS–PAGE of TwinStrep (TS)-tagged SsPepco following cleavage by SsAvs4. Band 1 corresponds to the cleaved monomer lacking the C-terminal fragment. (**B**) Mass spectrometry peptide coverage for band from (A).

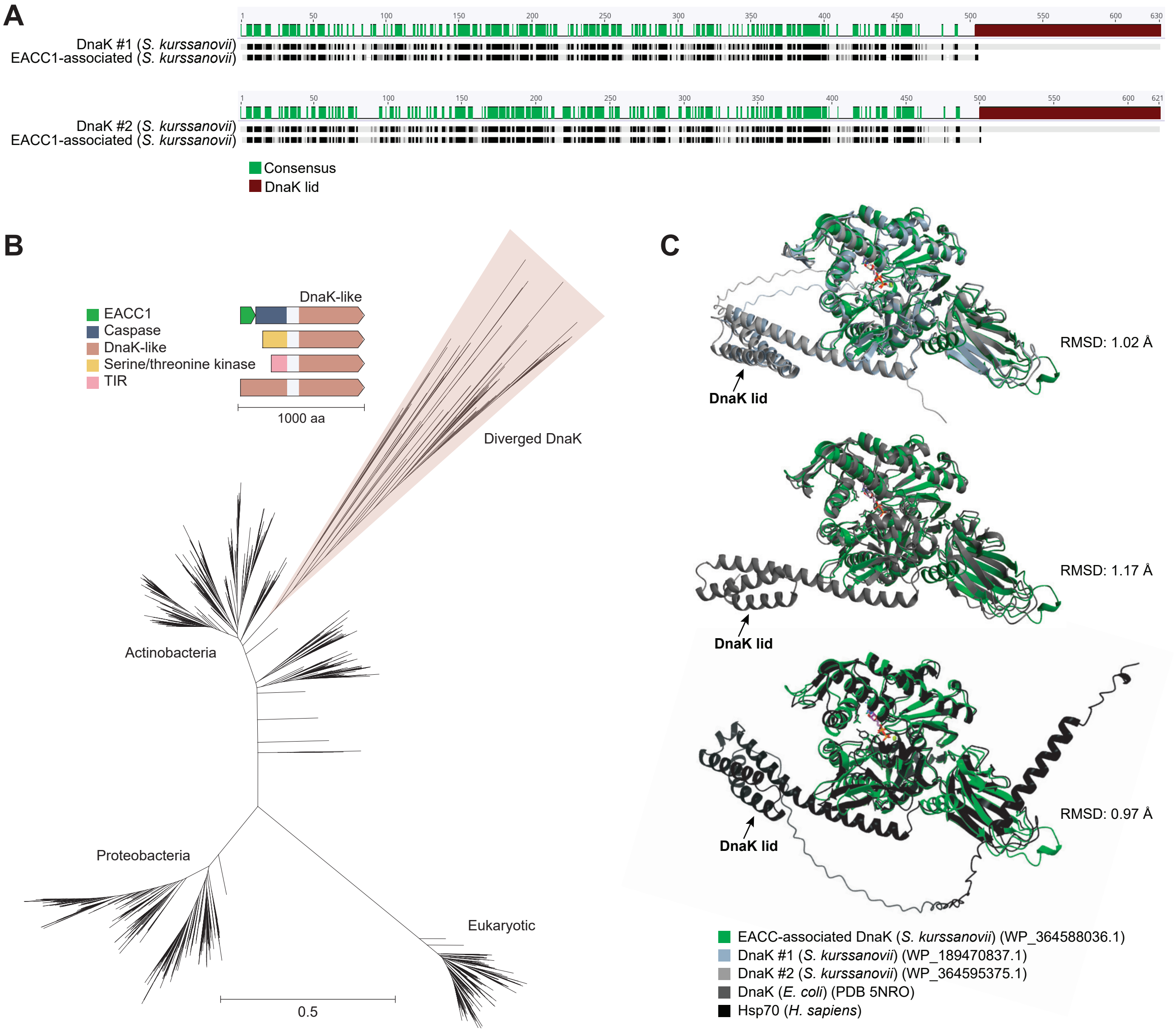


**Supplementary Figure 12. An EACC–caspase module fused to a DnaK-like chaperone (related to Figure 6).** (**A**) Pairwise alignment of EACC1 associated with a DnaK-like chaperone with each DnaK copy from *Streptomyces kurssanovii* NPDC049344. (**B**) Phylogenetic tree based on BLAST searches of the NCBI clustered_nr database using the EACC1-associated DnaK, *E. coli* DnaK, and human Hsp70. The core DnaK domains were aligned using MAFFT and clustered with MMseqs2 at 90% identity and 90% coverage (--min-seq-id 0.9 -c 0.9). The tree was constructed using FastTree (1,723 sequences). Divergent branches of DnaK containing N-terminal EACC1-caspase ($n$ = 66), serine/threonine kinase ($n$ = 19), DnaK ($n$ = 3), or TIR ($n$ = 1) domains all lack the C-terminal lid domain present in canonical DnaK chaperones. (**C**) Structural alignment of the EACC1-associated DnaK model with each DnaK copy in the *S. kurssanovii* genome, the *E. coli* DnaK structure (PDB: 5NRO), and a model of human Hsp70. All models were generated using AlphaFold3 with ATP and an $Mg^{2+}$ ion.

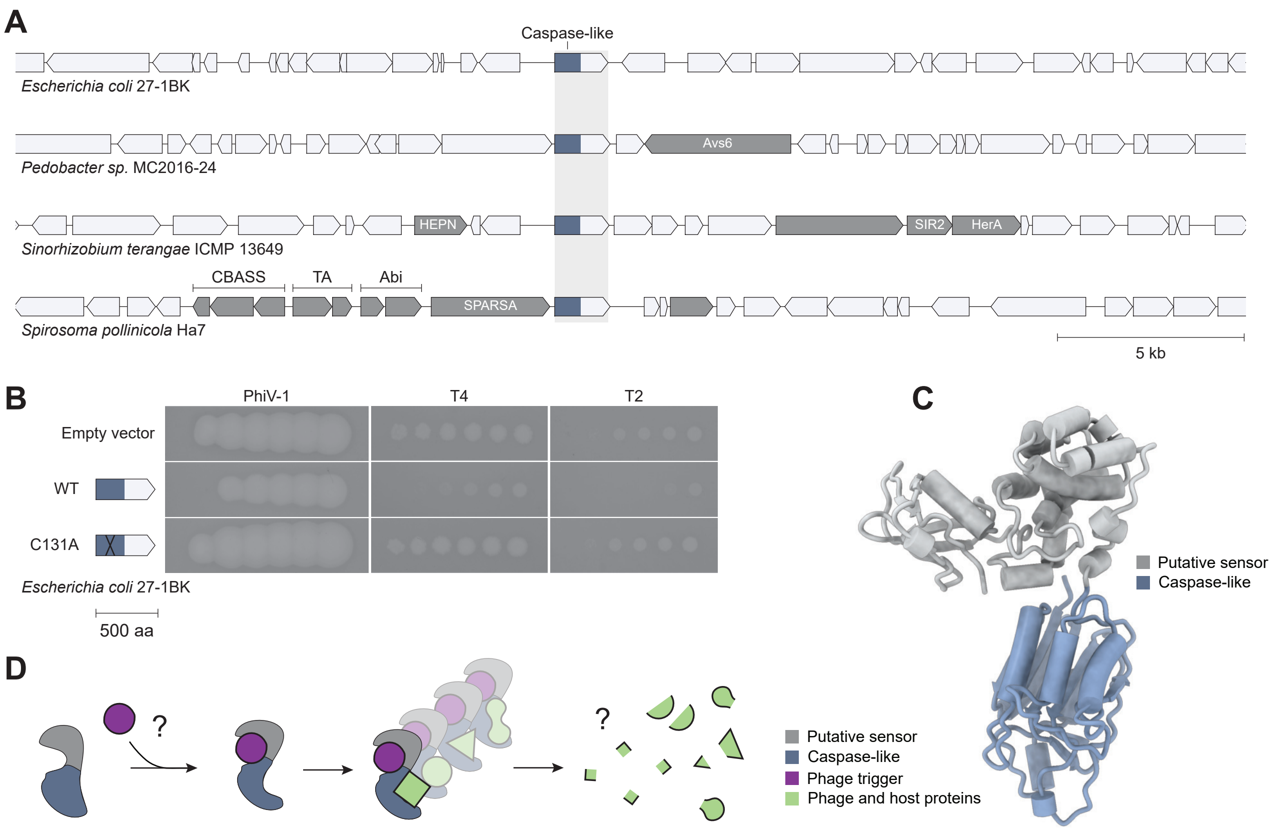


**Supplementary Figure 13. A caspase-like protease without an associated death effector confers phage defense.** (**A**) Representative genomic neighborhoods of a novel caspase-like defense protease without an associated death effector. (**B**) Heterologous reconstitution of the caspase-like protease from *E. coli* 27-1BK in *E. coli* K-12. Plaque assay spots show 10-fold serial dilutions of phages PhiV-1, T4, and T2 on strains expressing the wild-type or catalytic mutant (C131A) protease. (**C**) AlphaFold3 structural model of the caspase-like protease. (**D**) Schematic illustrating a proposed defense mechanism via general protease activity.
